# Rapid and repeated evolution of myosin copy number in threespine stickleback

**DOI:** 10.64898/2025.12.22.696110

**Authors:** Alyssa M. Yoxsimer, Rhea R. Daugherty, Emily E. Hare, Yingguang Frank Chan, Felicity C. Jones, Garrett A. Roberts Kingman, Emma G. Offenberg, Timothy R. Howes, Haili Zhang, Alex A. Pollen, Shannon D. Brady, Kathleen T. Xie, Heidi I. Chen, Craig B. Lowe, Eric H. Au, Jane Grimwood, Jeremy Schmutz, Richard M. Myers, Dolph Schluter, David C. Heins, Miguel L. Reyes, John A. Baker, Bjarni Jónsson, Thomas E. Reimchen, Michael A. Bell, David M. Kingsley

## Abstract

Copy number variants at genomic loci evolve at a high rate, are linked to many different diseases, and play a role in adaptive evolution in humans and other organisms. Here we show that stickleback fish from freshwater environments have rapidly and repeatedly evolved an expanded number of copies of a gene family involved in muscle development, *Myosin Heavy Chain 3 Cluster C* (*MYH3C*), compared to marine populations. Differences in copy number between marine and freshwater fish are maintained even in the presence of gene flow, suggesting that *MYH3C* changes represent adaptive divergence between ecotypes. Copy number expansion occurs by tandem duplication of *MYH3C* coding and regulatory regions on the stickleback sex chromosome. We identify a muscle regulatory enhancer within the expanded *MYH3C* region and show that elevated copy number is associated with developmental and tissue-specific increases in corresponding mRNA expression levels in skeletal muscle. Common *MYH3C* clusters include 3-, 4-, 5-, and 6-copy variants that likely evolved through a combination of microhomology-mediated break repair and non-allelic homologous recombination. Our results provide a new example of copy number changes in a wild species and identify CNVs as potential “hotspots” of repeated adaptive evolution.

## INTRODUCTION

Copy number variations (CNVs) are the most abundant type of variation in vertebrate genomes by total base pairs affected^1^ and make up approximately 12% of the human genome^2^. CNVs can change the structure of genes at breakpoints or alter levels of expression of dosage-sensitive genes^3^. In humans, these changes have been linked to many common traits and diseases, including red-green color blindness^4^, Charcot-Marie-Tooth disease^5^, schizophrenia^6^, autism^7^, Crohn’s disease^8^, and many others^3^. CNVs may also play an important role in adaptive evolution, including the evolution of herbicide resistance in plants, drug resistance in insects, and the adaptation of humans and other animals to new diets^9–13^. However, identifying CNVs that are adaptive rather than deleterious is challenging, and many known CNVs may have become common either by drift or inefficient purifying selection in duplicated gene sequences^14,15^.

Repeated evolution in similar environments provides powerful biological evidence that specific traits or genomic changes are positively selected. If very similar traits evolve independently in multiple lineages in response to similar ecological conditions, the recurrent changes have likely conferred an advantage^16,17^. Threespine stickleback (*Gasterosteus aculeatus*) are a particularly favorable system for studying repeated evolution because these fish have undergone recent, wide-scale, and replicated evolution following northern hemisphere glacial retreat within the past 20,000 years. Migratory marine (i.e., anadromous) stickleback colonized tens of thousands of new freshwater lakes and streams generated since the end of the last ice age and have repeatedly evolved similar changes in the new freshwater environments^18^. For example, marine stickleback are highly armored endurance swimmers that have long, streamlined bodies useful for long-range migration during annual movements between ocean foraging and coastal or freshwater breeding areas. By contrast, freshwater stickleback generally have reduced armor, deeper bodies, and greater body flexibility and burst swimming speed^19,20^.

Genomic studies have begun to identify many loci that also show consistent sequence differentiation between stickleback from marine and freshwater environments. Early genomic studies focused on regions with consistent base pair divergence detected by sequencing of key trait loci^21,22^, or by surveying many different loci using SNP genotyping arrays^23^, RAD-tag sequencing^24^, or full genome sequencing^25,26^. More recent studies have identified dozens to thousands of genomic regions that also show variable copy number differences among different stickleback populations^27–29^. With a few exceptions^12,29^, most of these potentially interesting CNV regions have not yet been further studied.

Here we show that a tandem cluster of *myosin heavy chain 3* (*MYH3*) genes shows recurrent copy number differences in stickleback that are consistently correlated with environmental conditions. Increased copy number rapidly evolves when marine fish colonize or are introduced to freshwater habitats. The class II myosin heavy chain genes that encode the characteristic sarcomeric motor proteins of skeletal muscle are included among the multiple myosin genes found in many organisms^30–32^. We find that increased copy number of the stickleback sarcomeric *myosin heavy chain cluster C* genes (*MYH3C*) in freshwater ecotypes is due to local tandem duplication of both myosin coding and flanking regulatory sequences, leading to tissue-specific increased expression of *MYH3C* in skeletal muscle. Long-read sequencing confirms the structures of the expanded clusters, and suggests molecular mechanisms that contribute to the origin of variable myosin arrays in natural populations.

## RESULTS

### Myosin copy number increases are recurrent and rapid in freshwater stickleback populations

We have previously carried out genome-wide screens for genomic regions that show either repeated SNP divergence between marine and freshwater stickleback (EcoPeaks), rapid allele frequency changes when marine fish are experimentally introduced to freshwater habitats (TempoPeaks), or evidence of structural rearrangements when comparing marine artificial chromosomes (BACs) to the freshwater reference genome^25,26,33^. Comparing these different patterns revealed a cluster of four tandemly arranged stickleback *MYH3* genes on the sex Chromosome XIX of the reference genome (Bear Paw Lake, AK [BEPA]) in a region that shows a significant EcoPeak, a significant TempoPeak, and different physical sizes in BACs and the reference genome (Figure 1A). The third and fourth *MYH3* genes share greatest sequence identity with each other and occur within a larger region of ∼17.6 kb that is tandemly duplicated.

**Figure 1.**
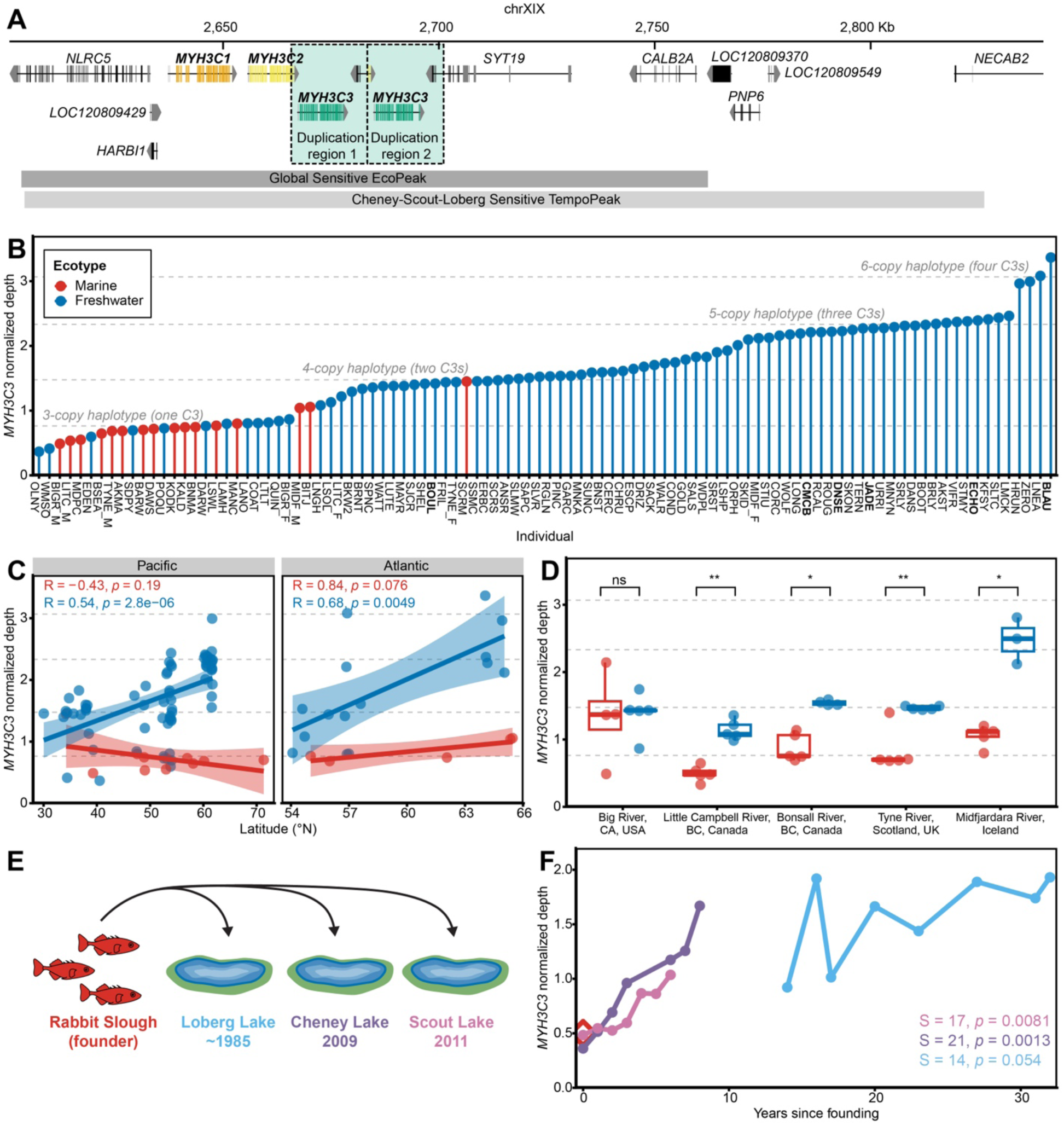
Myosin copy number increases are recurrent and rapid in freshwater stickleback populations. (A) ChrXIX genomic region in the freshwater stickleback reference genome (*stickleback v. 5*)^38^ that includes four tandemly arrayed myosin heavy chain genes: *MYH3C1* (orange), *MYH3C2* (yellow), and two duplicated copies of *MYH3C3* (green). NCBI gene annotations for nearby genes are shown in grey. The boundaries of the ∼17.6 kb duplicated regions (light green shading with dashed boxes) include full duplications of *MYH3C3* and partial duplications of *MYH3C2* and *SYT19*. The duplicated sequence falls within a region with repeated SNP divergence between global marine and freshwater stickleback populations (Global Sensitive EcoPeak; dark grey bar) and a region that shows rapid SNP allele frequency changes in three populations of marine fish introduced into freshwater lakes (Cheney-Scout-Loberg Sensitive TempoPeak; light grey bar) described by Roberts Kingman et al.^26^ (B) *MYH3C3* read depth normalized by mean autosomal read depth determined for female stickleback collected from different populations in the North American Pacific region (*n* = 10 marine, *n* = 66 freshwater) and Northern Europe (*n* = 5 marine, *n* = 15 freshwater). *MYH3C3* read depth is significantly higher in freshwater (blue) compared to marine (red) fish (two-sided Wilcoxon rank sum test, *p* = 9.83×10^-8^). Dashed lines indicate read depth calculated from simulated reads of different assembled 3- to 6-copy myosin haplotypes (one to four *MYH3C3* copies per haplotype; Figure 2). Bolded individual fish names indicate that a *MYH3C* region assembly was generated using another stickleback from the same population. See Table S1 and Roberts Kingman et al.^26^ for information about each sample. (C) Relationship between latitude and *MYH3C3* read depth for samples shown in panel B. *MYH3C3* read depth is positively correlated with Northern latitude in freshwater (blue) but not marine fish (red) in both the Pacific and Atlantic oceans (Pearson correlation). Lines show linear regressions with shaded 95% confidence intervals for each ecotype. (D) Box plots of *MYH3C3* read depth in five independent marine-stream systems from the Pacific and Atlantic basins where migratory marine (red) stickleback can meet and hybridize with freshwater resident (blue) stickleback. For all comparison groups *n* = 5, except for Big River marine (*n* = 4), Bonsall River freshwater (*n* = 4), and Midfjardara River freshwater (*n* = 3), In four of five systems, stream populations maintain significantly increased *MYH3C3* copy number compared to the marine counterpart in the same river (two-sided Wilcoxon rank sum test, ns = not significant, *p* > 0.05, \**p* ≤ 0.05, \*\**p* ≤ 0.01). (E) Independent introduction experiments were carried out in 2009 and 2011 by transplanting ∼3000 live marine stickleback from Rabbit Slough, AK into Cheney or Scout Lakes^43^. The first-generation progeny (0 years since founding) were recognized by size, and annual samples were collected through 2017. Loberg Lake was colonized by marine stickleback (most likely from nearby Rabbit Slough) between 1983 and 1988 after the lake was treated with rotenone to remove resident fish^44^. (F) *MYH3C3* read depth normalized to ChrXIX mean depth calculated from DNA sequencing of 39-200 pooled fish per sample. *MYH3C3* read depth increases significantly in Cheney (purple) and Scout (pink) Lakes in the first 6-8 years since introduction to freshwater (one-sided Mann-Kendall trend test). Loberg Lake (light blue) is already at a higher read depth at the first time point (14 years post founding) and has a positive but not significant upward trend thereafter. See also Figures S1-4.

Nomenclature and orthology relationships are difficult to assign in the multigene myosin heavy chain family, which has undergone multiple expansions, translocations, and gene conversions in different species^30–32^. However, the sequences of the stickleback ChrXIX *MYH3* genes, and the syntenic relationships of the surrounding genes, identify the ChrXIX cluster as the stickleback homolog of the “Cluster C” myosin heavy chain genes previously described in other percomorph fishes^30^. “Cluster C” genes: (1) belong to the *MYH3* family when aligned with known myosin heavy chain genes in humans^30,34^; (2) encode “fast” myosin heavy chains found in skeletal muscle^30,32^; and (3) are expressed in early developmental stages as well as in adult fish^35,36^. Basic Local Alignment Search Tool (BLAST)^37^ searching shows that a previously described myosin cDNA clone isolated from dissected adult trunk skeletal muscle of stickleback^32^ also has 100% identity across its 558 bp length to exons in the ChrXIX region, further confirming that *MYH3* Cluster C genes from the stickleback ChrXIX locus are spliced and expressed in adult skeletal muscle. Based on the sequences and synteny relationships with other fishes, we refer to the stickleback ChrXIX *MYH3-*related Cluster C genes as *MYH3C* genes, and we sequentially number the tandemly arranged genes in the cluster *MYH3C1* (*C1*), *MYH3C2* (*C2*), and *MYH3C3* (*C3)*; with the *C3* gene present in two copies in the freshwater reference genome^38^ (Figure 1A).

We tested for repeated evolution of *C3* copy number by calculating read depth from whole genome resequencing data of female marine and freshwater fish from representative populations in the Pacific and Atlantic Ocean basins previously used for global SNP divergence surveys^26^. We found that freshwater fish have a higher mean *C3* copy number compared to marine populations (two-sided Wilcoxon rank sum test based on one individual per population, n = 96 populations, *p* = 9.83×10^-8^; Figure 1B, Table S1). Comparing observed read depth to simulations, total copy number in the ChrXIX cluster haplotypes resembles the range expected from a minimum of three myosin genes in a cluster (*C1*, *C2*, plus a single *C3*) to a maximum of six myosin genes (*C1*, *C2*, plus four *C3*s). Haplotypes predicted to have one *C3* copy were most abundant in marine populations, while haplotypes with two to four *C3* copies were most abundant in freshwater populations. *C3* read depth was positively correlated with increasing latitude in freshwater stickleback in both the Pacific and Atlantic basins, suggesting higher copy numbers may be advantageous in northern freshwater environments (Figure 1C). Variable *C3* copy numbers could also be detected within several populations (Figure S1, Table S1).

To determine if increased *C3* copy number has likely evolved once or many times, we built a phylogenetic tree using 100,000 genome-wide neutral SNPs scored in the same stickleback shown in Figure 1B. Populations with increased *C3* copy numbers were widely distributed in the phylogenetic tree, and were interspersed with other populations showing lower copy numbers (Figure S2). Copy number changes thus appear to be evolving repeatedly in many different geographic locations, rather than in a particular subset of stickleback that are closely related by lineage.

To further explore possible selective pressures acting on myosin *C3* copy number variation, we tested whether the differences seen between populations are also observed within single rivers where migratory anadromous fish breed in sympatry with freshwater residents and form hybrid zones. Gene flow should homogenize most neutral genomic variation, except in genetic regions that are linked to sequences controlling ecotypic differences between the anadromous and freshwater forms or that interact epistatically with other such regions. Using the same methods and source dataset as the global survey^26^, we calculated *C3* read depth for three Pacific Ocean basin rivers (Big River, CA, USA, Little Campbell River, BC, Canada, and Bonsall River, BC, Canada) and two Atlantic Ocean basin rivers (Tyne River, Scotland, UK and Midfjardara River, Iceland) having well-characterized marine-freshwater hybrid zones^39–42^. In all rivers except for Big River (which occurs at the lowest latitude), the upstream freshwater stickleback show significantly higher *C3* copy numbers than downstream anadromous fish from the same river system (Figure 1D, Table S1). Thus, *C3* copy number differences are maintained even in the presence of gene flow.

Having observed greater *C3* copy number repeatedly in many post-glacial freshwater populations compared to marine stickleback, we tested whether we could also observe increases in copy number when marine populations were recently introduced into new freshwater habitats (Figure 1E). We transplanted marine fish from Rabbit Slough, AK (RABS) to Cheney and Scout Lakes, AK in 2009 and 2011, respectively^43^. Marine fish were also introduced to Loberg Lake, AK between 1983 and 1988 by unknown means^44^. *C3* read depth was calculated from sequenced pools of 39-200 fish from annual surveys of each lake after introduction^26^. Within the first six to eight years after founding, Cheney and Scout Lakes showed significant increases in *C3* copy number (Figure 1F). The Loberg Lake population is older and already showed a higher *C3* read depth than the presumptive marine founder population at the first collection time point (∼14 years post-founding), and continued to show an increasing trend in subsequent years. SNP allele frequencies previously determined for the *MYH3C*-spanning TempoPeak by Roberts Kingman et al. suggest that a higher copy number freshwater allele rose to a frequency of over 30% within six generations (Figure S3), with estimates of the selection coefficient *s* ranging from 0.182 to 0.574 depending on the method of calculation^26^. Together, these results indicate that changes in *C3* copy number are repeated across multiple marine-freshwater ecotypic pairs and also evolve rapidly following introduction of marine stickleback to new freshwater habitats.

### Myosin haplotypes

The high sequence identity of duplicated regions can make CNVs difficult to reveal in genome sequencing projects because duplicates are prone to dropout during assembly. To further investigate the structures of *MYH3C* gene clusters in marine and freshwater stickleback, we (1) reexamined previously published reference genomes, (2) carried out new *de novo* assemblies from our own long-read DNA sequencing of particular populations, and (3) sequenced multiple large-insert bacterial artificial chromosome (BAC) clones available for both the original freshwater stickleback reference genome (Bear Paw Lake, AK [BEPA]) and for a marine population (Salmon River, BC, Canada [SALR]; Table S2)^25,45^. Validation of each of these approaches involved mapping long reads to their corresponding assemblies and only retaining assemblies with even read coverage and no high-frequency sequence differences between mapped reads and the assembly in the *MYH3C* region. Together these approaches identified 3-copy haplotypes in two marine populations (RABS^46^, and SALR), 4-copy haplotypes in the F5 generation of a female Boulton Lake, BC, Canada x male Bodega Bay, CA cross (BOULxBDGB)^47^, and in University of Hull, UK, Cottingham Pond population (HULL), 5-copy haplotypes in six freshwater populations (BEPA, Echo Lake [ECHO], and Jade Lake [JADE], all from the Mat-Su Valley in AK; and Community Club Lake [CMCB], Denise Lake [DNSE], and Kalifonsky Lake [KFSY], all from the Kenai Peninsula in AK), and a freshwater 6-copy haplotype in Blautaver Lake, Iceland (BLAU; Figure 2A; Table S2). Every assembly had a single copy of *C1* and *C2*, with overall copy number variation coming from varying numbers of *C3* genes. Because we assembled a 5-copy haplotype from a BEPA fish using PacBio HiFi reads, we wondered whether the previous 4-copy count for the stickleback reference genome (derived from a BEPA fish)^25,38^ might be an underestimate (Figure 1A). We sequenced all seven BACs spanning *C3* duplications from the original BEPA fish used for reference genome assembly, and confirmed with individual long Nanopore reads that the reference fish was likely homozygous for a 5-copy haplotype (Figure S4).

**Figure 2.**
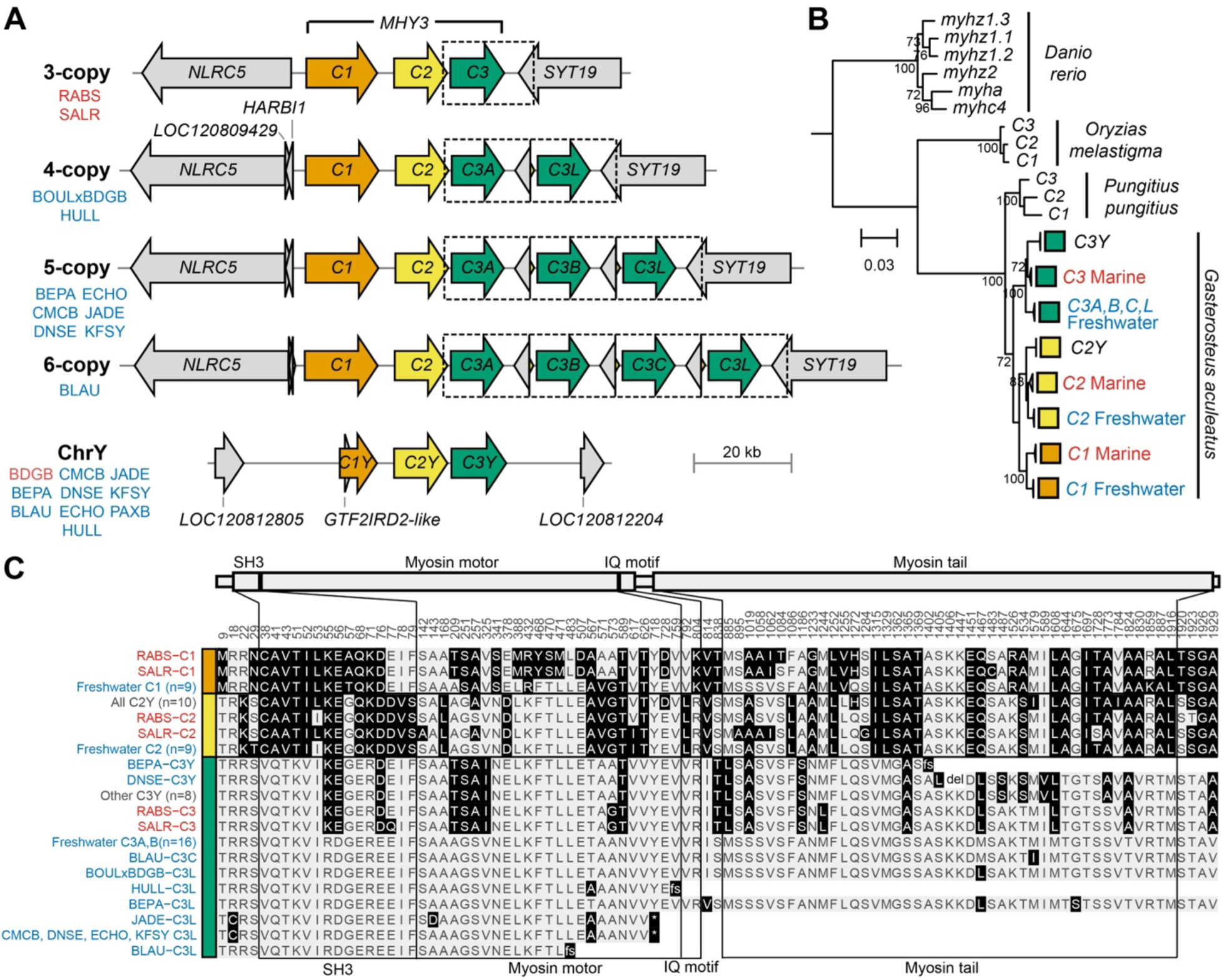
Myosin haplotypes and sequence variants from marine and freshwater stickleback. (A) Structure of *MYH3C* copy number haplotypes with one *C1* copy (orange), one *C2* copy (yellow), and one to four *C3* copies (green). All expansion haplotypes contain the same duplicated ∼17.6 kb region (dashed boxes) with breakpoints in *C2* and *SYT19* arranged in tandem. The 3-copy haplotypes come from marine (red) fish while 4-, 5-, and 6-copy haplotypes come from freshwater (blue) populations and show characteristic marine versus freshwater SNP divergence. The *C3* duplications occur exclusively on ChrXIX while a singular marine-like 3-copy haplotype with a pseudogenized *C1* is present on ChrY in all surveyed populations. *HARBI1* is also present between *NLRC5* and *C1* in freshwater 4- to 6-copy alleles but the first exon is deleted in BLAU and it is completely absent from marine 3-copy haplotypes. See Table S2 for sample abbreviations, source locations, and assembly information. (B) Phylogenetic tree based on coding region nucleotide sequences of threespine stickleback *MYH3C* copies and putative orthologs from *Danio rerio*, *Orzias melastigma*, and *Pungitius pungitius*. Bootstrap values of 270 are labeled below each node. Branch lengths are based on the number of inferred substitutions, indicated by the scale bar. *O. melastigma* and *P. pungitius MYH3C* copies were renamed *C1*, *C2*, and *C3* based on similar arrangement and location in a syntenic region to *G. aculeatus MYH3C*. All *MYH3C* copies form within-species clades. Within *G. aculeatus*, all *MYH3C* copies group first by copy (*C1*, *C2*, or *C3*), then by marine, freshwater or ChrY copies, with clades collapsed due to low confidence resolving relationships of the same *MYH3C-*copy-type and ecotype. (C) Myosin protein domains and amino acid differences found in different *MYH3C* copies and populations. Black highlights indicate an amino acid sequence difference relative to the most abundant protein sequence from freshwater C3A and C3B (light grey). DNSE C3Y contains a deletion of residues 1406-1447 (del). Several C3L or C3Y sequences contain premature stop codons caused by either nonsense mutations (*) or frameshift mutations (fs) which produce short (10-27 aa) stretches of different residues before reaching a stop codon (not shown). The SH3 domain is at 33-83 aa, the myosin motor domain is at 87-774 aa, the IQ motif is at 777-806 aa, and the myosin tail is at 842-1919 aa. See also Figures S4 and S5, and Table S3.

CNV can arise from either local or dispersed copies of a gene family^48^. In the current case, both the tandem gene assemblies and formal QTL mapping of *C3* copy number in a freshwater by marine genetic cross (BOULxBDGB) support *C3* copy number expansion exclusively at the *MYH3C* ChrXIX locus (Figure S5). Inspection of the myosin sequence assemblies shows that copy number expansions consist of one or more duplications of a similar tandemly arranged ∼17.6 kb region, which includes a full length *C3* gene copy and has breakpoint positions located in *C2* and the flanking gene, *SYT19*. Based on high similarity of sequences between expanded *C3* copies, we name the first *C3* copy *C3A* and the last *C3* copy *C3L (L* referring to *last)*. Any intermediate copies are named alphabetically (*C3B*, *C3C*) based on linear order in the cluster.

ChrXIX is a sex chromosome—the X chromosome in a recently evolved XY system^49^—so we also investigated ChrY *MYH3C* structure in assemblies derived from males. ChrY contains a marine-like 3-copy haplotype, although *C1* appears to be pseudogenized. The *MYH3C* region is located in the male-linked, non-pseudoautosomal stratum 2 of the Y chromosome, and shows no evident amino acid divergence between marine and freshwater stickleback populations (Figure 2)^50^.

To evaluate the relationships between *MYH3C* copies in stickleback and other fish outgroups, we constructed a phylogenetic tree based on *MYH3C* protein-coding nucleotide sequences. Putative *MYH3C* orthologs were identified in zebrafish (*Danio rerio*), which have two inverted clusters of three tandem myosin genes each, Indian medaka (*Orzias melastigma*), which have three tandem myosin genes in a region syntenic to *Gasterosteus*, and ninespine stickleback (*Pungititus pungitius*), which also have three syntenic, tandem myosin genes. While each species showed three or more *MYH3C* genes, the sequences of these myosins formed within-species rather than between-species groups. This tree topology suggests that either each lineage has independently expanded to three or more tandem myosin copies, or that extensive gene conversion has homogenized an ancestral three-gene group within each species since divergence^32^ (Figure 2B). Marine and freshwater stickleback ChrXIX *MYH3C* copies form distinct clades within *Gasterosteus*, with all *C3* expanded copies grouping together. This tree structure indicates an ancestral 3-copy myosin haplotype in threespine stickleback, then divergence of marine and freshwater haplotypes, followed by recent expansion of *C3* copies through additional duplications on the freshwater haplotype.

Gene duplicates may retain original functions, gain new functions (neofunctionalization), partition old functions (subfunctionalization), or lose functions^51^. To examine whether *MYH3C* copies show any amino acid changes that could indicate divergence in function, we aligned the protein sequences of all *Gasterosteus MYH3C* copies and extracted divergent amino acid positions. *C1*, *C2*, and *C3* each encode a 1931 amino acid (aa) protein, with 96.8-98.7% pairwise amino acid identity between full length C1, C2, and C3 proteins, and even higher 98.9-100% identity among the duplicated full length C3 proteins. There are 16, 2, and 18 aa changes in all freshwater compared to marine C1, C2, and C3 proteins, respectively; indicating that MYH3C protein sequence changes may contribute to differences in muscles between marine and freshwater fish (Figure 2C, Table S3). Marine-freshwater divergent amino acids in C3 are enriched in the SH3 domain (hypergeometric test, *p* = 0.01) and include three charged residues that change to a residue with the same charge. All freshwater C3A and C3B sequences are identical, suggesting that a dosage effect or modification of regulatory sequence, rather than a change in protein sequence, is most likely responsible for any phenotypic effect that these duplicated copies may confer. While some C3L sequences have amino acid changes relative to C3A, these changes are not consistent across populations. Furthermore, C3L in all assemblies except BOULxBDGB and BEPA has frameshift or non-sense mutations that truncate the protein sequence within the myosin motor domain, indicating that some copies may have become non-functional (Figure 2C).

### Myosin expression

In addition to changes in protein sequence, gene copies may diverge in function through changes in expression patterns. Co-expression of tandem duplicates may lead to dosage increases, but epigenetic silencing or sequence changes in regulatory regions may also reduce expression of expanded gene copies^52–54^. To test for possible *cis*-regulatory differences in developmental and tissue-specific patterns of expression between freshwater and marine CNV alleles, and across *MYH3C* copies, we crossed fish carrying 5-copy freshwater BEPA and 3-copy marine RABS haplotypes, and quantified allele-specific expression (ASE) in F1 hybrid females that carry both myosin clusters in the same *trans*-acting environment (Figure 3A). F1 hybrids were raised until approximately 1 month or 5 months post fertilization, at which point the whole body was collected from 1-month juveniles, and separate flank muscle, pectoral abductor muscle and pectoral adductor muscle were dissected from the 5-month young adults. We performed RNA-seq on six individuals per condition to quantify both total expression levels between *MYH3C* copies and ASE differences between the marine and freshwater alleles. Because there are very few exonic variants that distinguish the *C3A*, *C3B*, and *C3L* gene copies, and long reads are necessary to differentiate all three freshwater *C3* transcripts, we also performed PacBio Kinnex sequencing on two individuals per condition.

**Figure 3.**
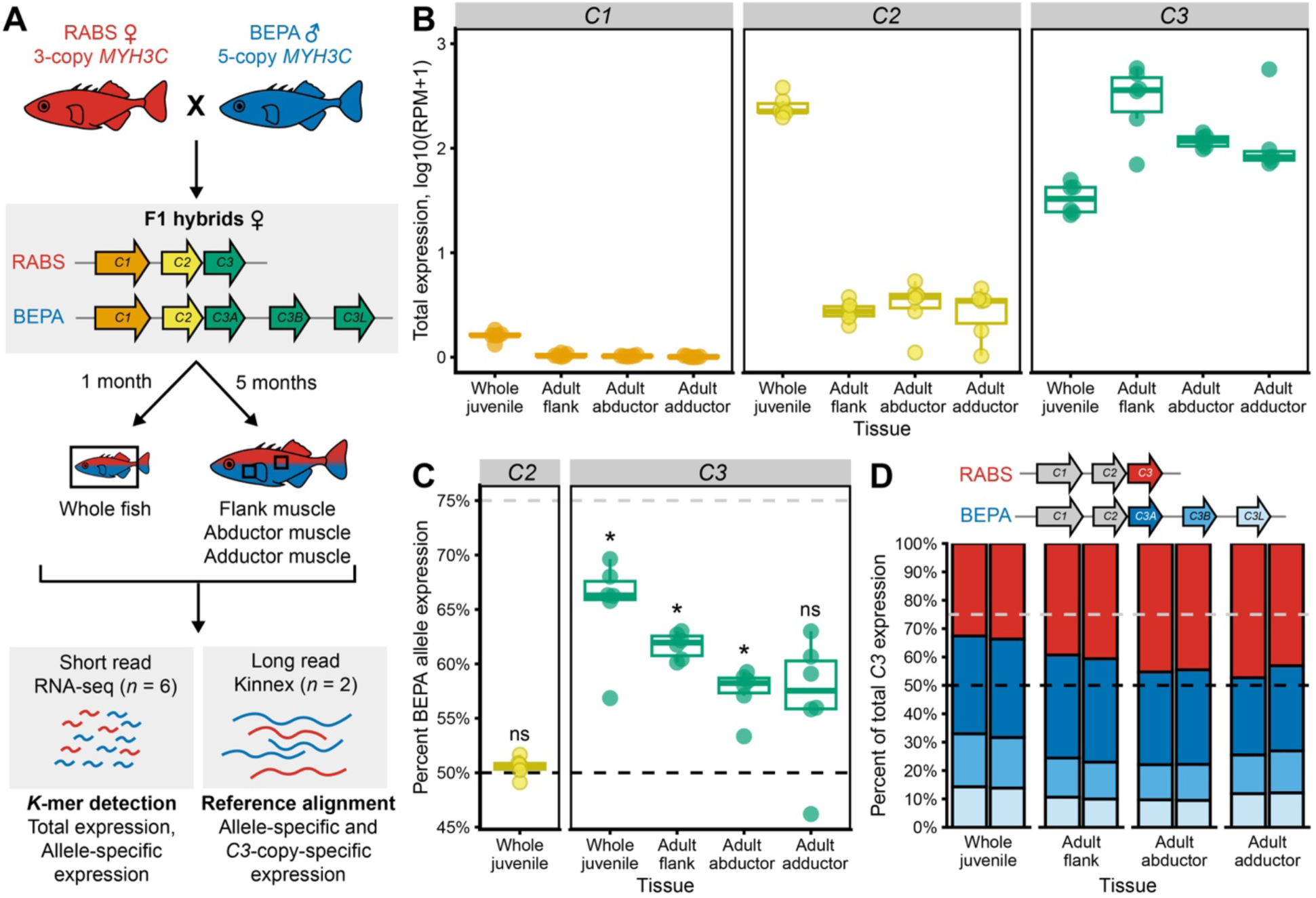
Expression of different *MYH3C* copies and alleles in developing stickleback. (A) F1 hybrids were generated by crossing two 3-copy marine RABS (red) females with a 5-copy freshwater BEPA (blue) male and raising offspring for either 1 or 5 months. RNA was extracted from whole juvenile fish (no caudal fin) from 1-month females, or from flank muscle, abductor pectoral muscle, and adductor pectoral muscle dissected from 5-month young adult females. *K*-mers were used to detect total expression or allele-specific expression (ASE) from RNA-seq reads (*n* = 6 per condition; Table S4). Alignment of Kinnex reads (*n* = 2 per condition) to a marine 3-copy and freshwater 5-copy composite genome was used to identify relative expression of each *C3* copy. For simplicity, the small fragments of the *C2* and *SYT19* genes are not diagrammed in the RABS and BEPA haplotypes. (B) Box plots showing the relative expression levels of *MYH3C* copies in each tissue based on seven sets of *MYH3C* copy expression (MCE) 27-mers. Reads per million (RPM) was calculated by dividing the number of paired reads detected by each *k*-mer by the total number of paired reads from each sample. *C1* (orange) has no to low expression in all tissues surveyed. *C2* (yellow) is highly expressed in juvenile whole fish. *C3* (green) is expressed most highly in adult flank muscle, but is highly expressed in all tissues surveyed. (C) Box plots showing allele-specific expression of the BEPA vs. RABS alleles based on ASE *k*-mer sets (four *C2* sets and nine *C3* sets) for each *MYH3C* copy. Deviation from an equal ratio of expression from the BEPA and RABS alleles (50% BEPA allele expression, black dashed line) was evaluated using a one-sample Wilcoxon rank sum test (mu = 0.5; ns = not significant, *p* > 0.05; \**p* < 0.05). Expression from the three combined *C3* copies of the BEPA allele is higher than expression from the single RABS *C3* gene in juvenile whole fish and adult flank and abductor muscle but not fully proportional to copy number (75% expected BEPA allele expression, grey dashed line). *C2* shows no ASE. ASE was not calculated for all *C1* tissues and *C2* adult tissues due to low expression. (D) Stacked bar plots representing the relative expression levels of *C3* copies from RABS (red) and BEPA (*C3A*, dark blue; *C3B*, medium blue; *C3L*, light blue) determined by mapped Kinnex reads. All duplicated *C3* copies are expressed, *C3A* is expressed at approximately twice the level of *C3B* and *C3L*, and the summed expression of all freshwater *C3* copies falls between 50% and 75% of total expression (lower and upper dotted lines), as expected for increased but not fully proportional expression from the expanded freshwater haplotype. See also Figure S6 and Table S4.

In order to minimize the impact of reference bias and mis-mapping on RNA-seq quantification of highly similar *MYH3C* copies, we designed two sets of *k*-mers based on alignment of BEPA and RABS genomic assembly sequences: (1) *MYH3C* copy expression (MCE) *k*-mers which detect equivalent positions in all *MYH3C* copies but distinguish *C1* from *C2* from *C3* (used for total expression levels) and (2) ASE *k*-mers that possess at least one variant that uniquely identifies one *MYH3C* copy (i.e. *C3* from *C1* and *C2*) and at least one variant that distinguishes the RABS and BEPA alleles. We screened paired reads for perfect *k*-mer matches, then normalized expression either to total reads (MCE) or calculated the proportion of BEPA reads from the sum of BEPA and RABS reads (ASE), and averaged the results across *k*-mer sets.

We found that *C1* had no to very low expression in all tissues surveyed, *C2* was very highly expressed in the juveniles, and *C3* was highly expressed in all tissues, but especially in young adult flank muscle (Figure 3B). Because the whole body was sampled at the 1-month stage, it is unclear whether the lower expression of *C3* in 1-month juveniles is due to a ubiquitous low level of expression in skeletal muscle or expression is limited to a subset of anatomical sites. *C3* showed significantly higher ASE of the BEPA (freshwater) allele both in juveniles and in adult flank and abductor muscle, as might be predicted from the increased copy number of *C3* in the freshwater BEPA vs. RABS marine myosin haplotypes. However, the 53-70% expression of the BEPA allele was not as high as the predicted expression (75%) if expression was directly proportional to the number of genomic copies (Figure 3C).

To determine what proportion of *C3* expression derives from each of the different duplicated gene copies, we aligned full length non-concatemer (FLNC) Kinnex reads to a hybrid BEPA x RABS reference assembly and counted the number of mapped reads fully spanning *C3*-copy informative variants. We found that all three BEPA *C3* copies were expressed, though *C3A* was expressed at approximately twice the level of *C3B* or *C3L* (Figure 3D). The relative proportions of *C3A*, *B*, and *L* remained similar across developmental time points and tissues.

To assess whether temperature may affect the expression of different *MYH3C* copies, we also exposed siblings of the BEPA x RABS F1 hybrids described above to cold temperatures (3°C vs standard 18°C). Because of limited numbers of animals, we were underpowered to detect ASE differences in adult 3°C fish (*n* = 5). We did observe a few temperature-specific effects on expression (Figure S6), but longer acclimation time and increased sample numbers are likely necessary for more robust detection of temperature effects.

### The duplicated C2-C3 intergenic region contains a muscle enhancer

In addition to the *C3* gene body, the ∼17.6 kb duplicated region includes non-coding regions between *C2* and *C3*, and *C3* and *SYT19* (Figure 2A). We hypothesized that these regions may include regulatory sequences relevant to *C3* function. To test this, we cloned a single copy of the ∼2.6-2.8 kb region between *C2* (exon 35) and *C3* (exon 3) upstream of a minimal promoter and GFP (Figure 4A). Fertilized stickleback eggs were injected with constructs derived from either a marine SALR fish (*n* = 17) or a freshwater BEPA fish (*n* = 21). Transgenic embryos showed GFP expression specifically in skeletal muscle fibers starting as soon as 13 days post fertilization (dpf) and continuing past 3 months of age (Figure 4B). This pattern is consistent with the known larval and skeletal muscle expression patterns of *MYH3C* genes previously reported in other percomorph fishes^35,36^. To control for possible background expression arising from the Hsp70 promoter, we also injected a basal promoter-GFP construct that lacked the *C2*-*C3* intergenic region (*n* = 53). The SALR and BEPA constructs drove significantly more GFP expression in developing muscle fibers than the empty control vector and no difference was observed between the SALR and BEPA constructs (Figure 4C). The *C3* duplication events thus included regulatory sequences that drive expression in stickleback skeletal muscle.

**Figure 4.**
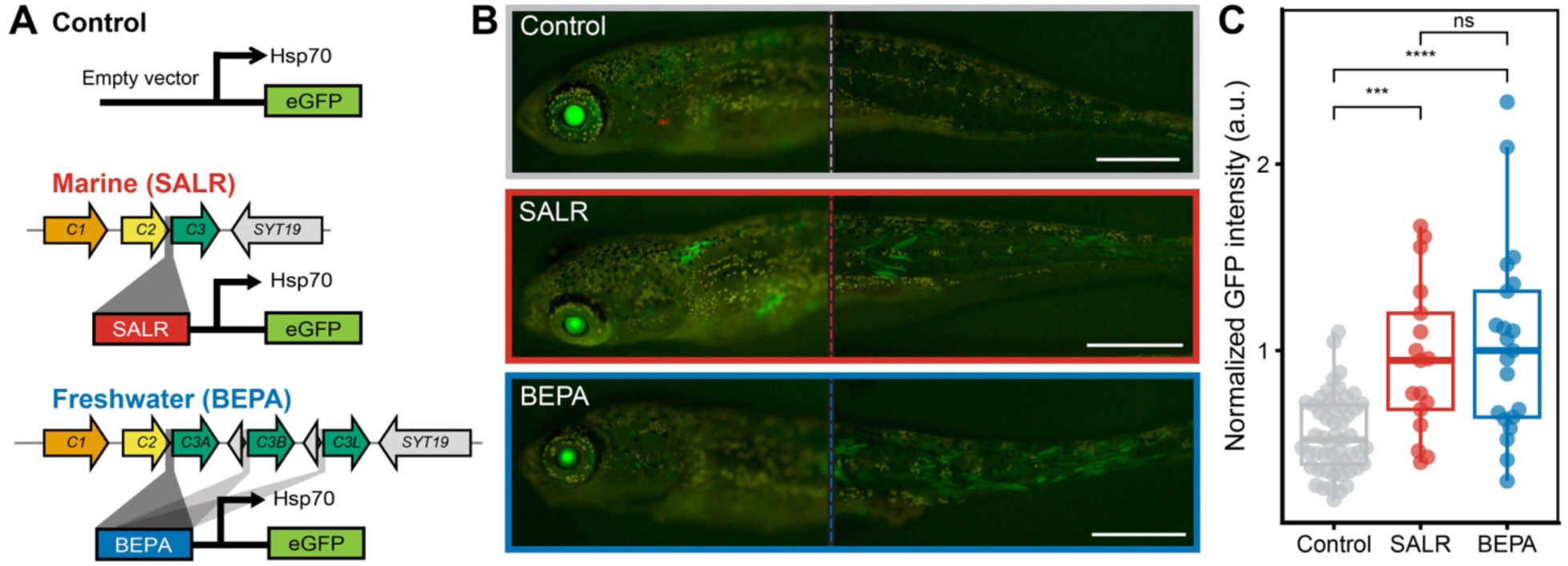
The duplicated *C2*-*C3* intergenic region contains a muscle enhancer. (A) Schematic of the negative control empty vector, SALR, and BEPA constructs used to generate transgenic fish. The SALR (red) and BEPA (blue) putative regulatory regions, which include the intergenic region between *C2* and *C3* (dark grey highlight), were inserted upstream of a minimal promoter (Hsp70) driving green fluorescent protein (GFP). The candidate regulatory intergenic region is present in one copy in the marine SALR 3-copy haplotype and three copies in the freshwater BEPA 5-copy haplotype—upstream of *C3A*, *C3B*, and *C3L*. For the expression constructs, a single copy of either the SALR or BEPA sequences was cloned and tested. (B) Representative images of GFP expression in transgenic stickleback. The SALR and BEPA constructs drive expression in the lens of the eye (left image) and in body skeletal muscle fibers of 3-week old larval fish (right image). Muscle expression begins at about 13 dpf and continues past 3 months of age (not shown). The control empty vector construct drives expression predominantly in the lens, a known site of activity of the Hsp70 promoter^84^. Scale bar, 1mm. (C) Box plots of normalized GFP intensity from multiple independent 3-week old transgenic fish carrying either the control empty vector (*n* = 53), SALR (*n* = 17), or BEPA (*n* = 21) constructs. The SALR and BEPA constructs drive significantly more expression in skeletal muscle than the control vector (two-sided Wilcoxon rank sum test of normalized flank muscle/eye intensity scores. Ns = not significant, *p* > 0.05, \*\*\**p* ≤ 0.001, \*\*\*\**p* ≤ 0.0001)

### Mechanisms of MYH3C copy number expansion

Gene duplications can arise recurrently or non-recurrently through diverse mechanisms including recombination-based, DNA replication-based, reverse transcription-based, and transposable element (TE)-mediated mechanisms. Two main mechanisms for change in gene copy number are non-allelic homologous recombination (NAHR) and microhomology-mediated break-induced replication (MMBIR)^55^. NAHR, the most common source of recurrent copy number changes, occurs when repeated sequences mispair during meiosis, resulting in unequal crossing over and duplication, deletion, or inversion^56^. By contrast, MMBIR does not rely on extended regions of homology and instead primes replication at stalled forks (typically after a double-strand break) using short 2-20 bp microhomology in nearby sequences^57^. Consequently, MMBIR typically results in variable duplications, deletions, or complex rearrangements depending on the position of breakpoint microhomology sites.

To determine whether *MYH3C* expansions show sequence signatures consistent with particular mutation mechanisms, we investigated the breakpoints and patterns of variation within expanded copies. When we compared the 4-copy freshwater haplotype to the 3-copy marine haplotype, we found that breakpoints in *C2* and *SYT19* were marked by 4-8 bp of microhomology in otherwise non-homologous regions (Figure 5A). These signatures are consistent with the MMBIR mechanism, where a break in *SYT19* stalls replication and replication is primed using shared microhomology found in *C2*. The resulting *SYT19*/*C2* junction between *C3* duplicates is present in all 4- to 6-copy haplotypes surveyed (Figure 2A), suggesting that a single, non-recurrent MMBIR event likely produced the initial 4-copy freshwater myosin haplotype before further expansion to five or six copies.

**Figure 5.**
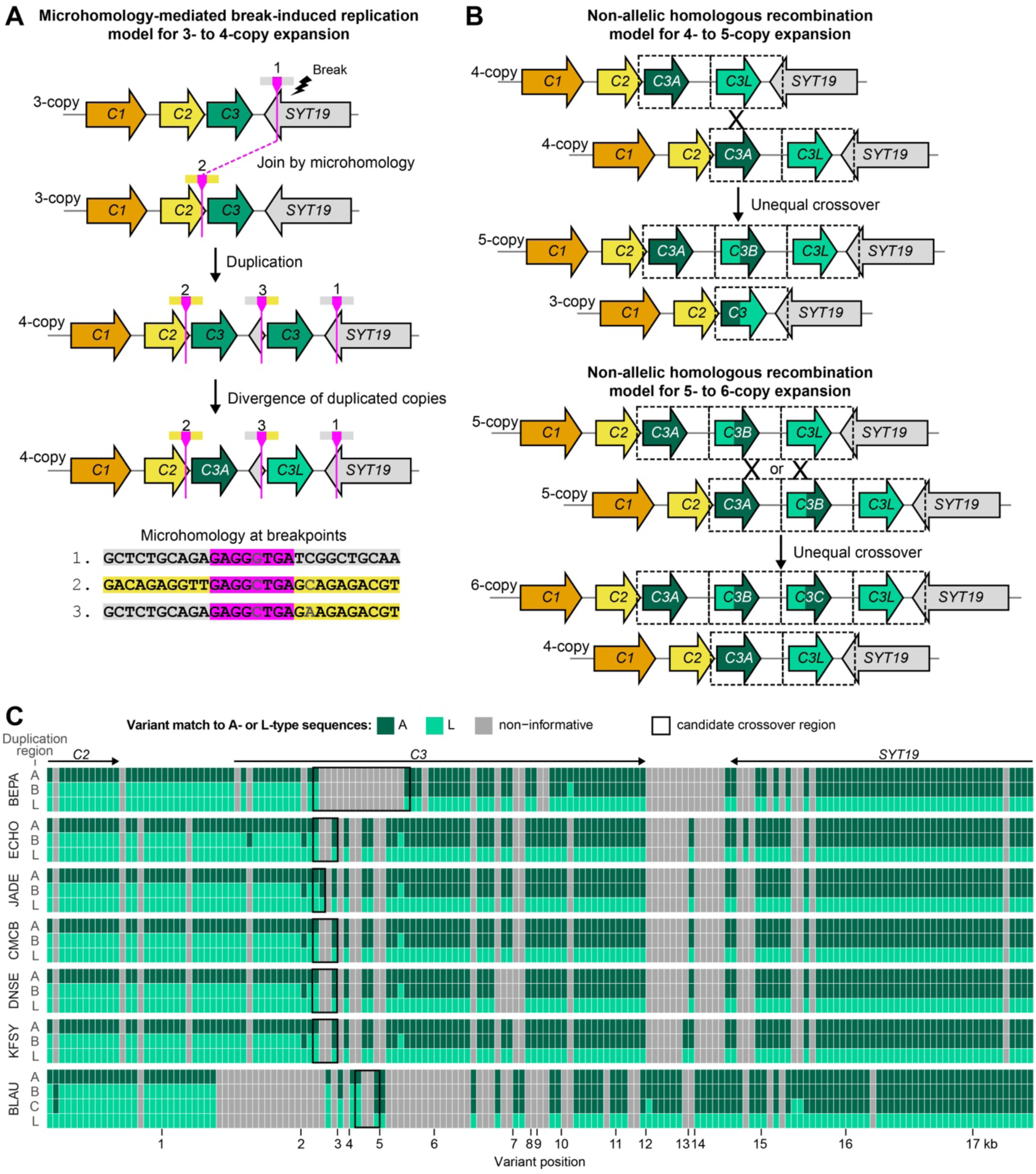
Mechanisms of *MYH3C* copy number expansion. (A) Hypothesized mechanism for 3- to 4-copy expansion via microhomology break-induced replication (MMBIR). During replication, a double strand break occurs near the *SYT19* breakpoint (1, grey) of a 3-copy haplotype, followed by 5’ to 3’ resection. Annealing to the sister chromatid or homologue using shared microhomology (fuchsia) within *C2* (2, yellow) restarts replication and produces a duplication with an intermediate breakpoint (3) that fuses the *SYT19* and *C2* sequences. Perfect microhomology of 4 bp and imperfect microhomology of 8 bp occurs at the breakpoints in all haplotypes (BOULxBDGB sequence shown here). (B) Hypothesized mechanism for 4- to 5-copy expansion and 5- to 6-copy expansion via non-allelic homologous recombination (NAHR). Inter-chromosomal or inter-chromatid recombination between two misaligned 4-copy haplotypes (*C3A* with the *C3L* duplicated region) could produce reciprocal duplication (5-copy) and deletion (3-copy) products (top). The new intermediate duplication gene would be more similar to the L region at the 5’ end and the A region at the 3’ end. If inter-chromosomal or inter-chromatid recombination occurred between two 5-copy haplotypes where *C3A* and *C3B* or *C3B* and *C3L* were paired, 6-copy duplication and 4-copy deletion products would be generated (bottom). Candidate recombination sites are indicated with an X but could theoretically occur anywhere along the ∼17.6 kb duplication regions (dashed box). For simplicity, the small fragments of the *C2* and *SYT19* genes are not diagrammed in all duplication boxes. (C) Nucleotide alignments of full ∼17.6 kb duplication regions from all 5 and 6-copy assemblies. Each colored box represents a divergent single nucleotide variant (alignment position indicated on the X-axis) that distinguishes the *C3A* duplication region (A; dark green) from the *C3L* duplication region (L; light green). Uninformative positions (grey) within assemblies are not shown (i.e. A, B, and L are identical or a variant is only found in B). Intermediate duplication regions (B, C) show the expected arrangement predicted by the NAHR model with the L-like sequence at the 5’ end and the A-like sequence at the 3’ end. Small regions that deviate from this pattern are likely due to gene conversion. The position of transition from the L to A allele (black box) indicates where a crossover event may have occurred. The candidate crossover site in BLAU (4,246-4,730 bp) differs from JADE (2,451-2,596 bp) and ECHO, CMCB, DNSE, and KFSY (2,451-2,932 bp) suggesting that expansion from a 4- to 5-copy haplotype may have occurred at least twice independently.

Once expanded to a 4-copy haplotype, the *C3A* and *C3L* tandem duplicates contain long stretches of highly similar sequence that can provide the substrate for NAHR^57^. Mispairing of *C3A* with *C3L* and unequal crossover would result in reciprocal deletion (3-copy) and duplication (5-copy) products that are predicted to have L-like sequence at the 5’ end and A-like sequence at the 3’ end of a new intermediately positioned *C3B* in the 5-copy haplotype (Figure 5B). With additional homology from a 5-copy gene cluster, further expansions through NAHR to six or more copies would be possible and could explain the presence of the 6-copy BLAU haplotype. To identify whether the intermediate *C3* copies found in the 5- and 6-copy haplotypes show the characteristic L/A junction pattern and identify candidate crossover sites, we aligned the ∼17.6 kb full duplication regions from all 5- and 6-copy assemblies. Within each assembly’s duplicated regions, we extracted positions where the *C3A*-containing duplication region (A) diverged from the *C3L*-containing duplication region (L) and then evaluated whether each position in the intermediate duplication regions (B,C) matched A or L. We found that the intermediate gene copies showed the characteristic L/A arrangement consistent with the expected NAHR duplication products (Figure 5C). We also found that while ECHO, JADE, CMCB, DNSE, and KFSY seem to share the same crossover site, the BLAU intermediate copies have a different putative crossover site, suggesting that 4- to 5-copy expansion has occurred independently in some populations.

## DISCUSSION

Our data provide compelling evidence that *MYH3C* copy number is repeatedly selected in threespine stickleback, with higher copy numbers evolving over and over again in freshwater populations compared with the ancestral marine state. These ecotype differences in copy number are maintained even in the presence of gene flow between marine and stream populations breeding within the same rivers, which should otherwise homogenize most neutral genomic regions. Increases in copy number are also seen in real time within a few generations after marine fish are transplanted into lakes, and the direction of these changes is concordant with the direction of differences present in many long-extant freshwater populations established at the end of the ice age. Molecular studies show that the expanded region contains an enhancer expressed in muscle, and that higher genomic copy number results in developmental and tissue-specific increases in *C3* gene expression. Finally, *MYH3C* CNVs likely emerged through multiple mutational events with a single original expansion from a 3- to 4-copy haplotype, followed by multiple expansions to 5- and 6-copy haplotypes.

Although *MYH3C* provides an interesting new example of genomic copy number changes consistently associated with particular environmental conditions, the selective advantages conferred by the changes in *MYH3C* still require additional study. The different members of the ancestral marine cluster of *C1*, *C2*, and *C3* show clear developmental and tissue-specific expression patterns as well as amino acid changes that suggest divergent specialized functions, as has been observed in “Cluster C” myosins of other percomorphs^35,36^. In addition, C1, C2, and C3 each show some amino acid changes that have diverged between marine and freshwater alleles. Marine-freshwater divergent amino acids in C3 are enriched in the SH3 domain, which facilitates interaction of the light chain with actin to modify the cycling kinetics of the myosin motor^58^. Three-dimensional modeling with AlphaFold^59^ suggests marine-freshwater amino acid differences occur at exposed sites on the heavy chain surface that could affect myosin interchain interactions, but additional functional studies will be required to measure the possible effects of these changes on myosin motor activity.

In contrast to the numerous amino acid differences between marine and freshwater C1, C2, and C3 molecules, the multiple expanded copies of C3 found in some freshwater populations show either no amino acid differences from each other (C3A and C3B), or a few scattered amino acid changes (C3L).

For this reason, we think the expanded freshwater haplotypes are more likely to act by conferring changes in myosin expression levels or patterns, rather than altered protein functions. Our expression studies using BEPA fish indicate that the expanded *C3* copies are all expressed, but with *C3B* and *C3L* at lower levels than *C3A*. The expression differences between the different C3 expanded copies may be due to shared sequence changes at the 5’ end of *C3B* and *C3L* (Figure 5C) that decrease the activity of their promoters, or due to the tendency of newly duplicated genes to be methylated^53^. As *MYH3C*s are highly expressed members of a larger protein complex, they may face strong selection on dosage due to energetic and stoichiometric constraints. Nonetheless, we do observe significantly higher overall expression of *C3* copies from the expanded freshwater haplotypes, though this dosage effect is not strictly proportional to copy number and appears to be specific to certain developmental time points and tissues.

Quantitative changes in the expression levels of myosin isoforms may affect biochemical properties, contraction speeds, or temperature optima of muscle^62–65^. In contrast to the superior endurance swimming of marine stickleback, freshwater stickleback exhibit better burst swimming, achieving higher maximum velocity and distance traveled in swimming performance tests^19,20^.While endurance swimming relies on the primarily red (slow-twitch, aerobic) and pink muscle fibers of the pectoral muscles, burst swimming is powered through the white (fast-twitch, anaerobic) fibers in flank musclature^19,66,67^. Based on the higher dosage effect seen in the juvenile body and adult flank muscle, and identification of a flank muscle enhancer, we hypothesize that the fast-twitch *MYH3C* duplicated copies may contribute to altered burst swimming speeds in freshwater fish. We have not yet been able to verify this prediction in limited behavioral experiments, but this idea could be further tested in the future with larger numbers of animals carrying contrasting myosin haplotypes on an otherwise uniform genetic background.

Particular myosin isoforms are also differentially expressed at different temperatures, and changes in myosin levels and isoforms may be a major mechanism to maintain muscle function in varying environments^30,62,68^. Many poikilothermic organisms have larger myosin gene repertoires than homeothermic organisms, and myosin gene expansion has previously been proposed as a convergent macroevolutionary mechanism used by poikilothermic animals to maintain muscle performance under a range of environmental temperatures^30^. We note that freshwater stickleback live for extended periods of the year at temperatures substantially higher and lower than the temperatures found in the ocean, and we observe that *MYH3C* copy number increases with latitude in freshwater populations. It is possible that expanded *MYH3C* copy numbers may be recurrently selected to maintain swimming performance under diverse conditions of temperature and oxygen concentrations in variable freshwater habitats, or as an adaptation to other factors that covary with latitude, such as season lengths that could influence developmental expression timing.

We also note that *MYH3C* variants are not the only differences between marine and freshwater fish in this ChrXIX genomic region. The previously reported EcoPeak on ChrXIX includes single full-length copies of several other linked genes including *SYT19*, *CALB2A*, most of *NLRC5*, and *HARBI1*. *SYT19* encodes a member of the synaptotagmin protein family, which act as calcium sensor proteins during vesicular transport and exocytosis in neuronal and endocrine tissues^69,70^. *CALB2A* encodes calbindin 2a, which is involved in neural development and motor pathways in fish^71^. *NLRC5* encodes a NOD-like receptor that plays an important role in regulation of both the innate and adaptive immune system during host responses to bacterial and viral pathogens^72–74^. *HARBI1* is anciently derived (450-500 million years) from a Harbringer transposase^75^ and may be involved in innate immunity and myosin regulation^76^. Interestingly, *HARBI1* is found only in the freshwater *MYH3C* haplotypes (although BLAU carries a deletion of exon 1). Future studies should investigate the individual contributions of both the tandemly duplicated *MYH3C* genes, and the other nearby linked genes in the surrounding EcoPeak region.

Because of the structural differences found between the X (ChrXIX) and Y versions of the *MYH3C* cluster, we note that XY male stickleback will carry fewer functional copies of *MYH3C* genes than XX females. This sexual dimorphism in *MYH3C* copy number is predicted to be larger in freshwater than marine stickleback because of the expanded number of myosins on the freshwater X-encoded myosin haplotype, and even larger in freshwater fish from northern latitudes that harbor even higher copy numbers on their X-linked myosin haplotype. The myosin copy number differences we describe here could thus contribute to general sexual dimorphism in muscle phenotypes between males and females, to sexual dimorphism that is more prominent in some populations than others, and to sexual dimorphism in physically linked non-myosin functions (for example, dosage of *HARBl1*). Sexual dimorphism is a prominent feature of many stickleback populations, and future studies can examine whether structural differences in the ChrXIX cluster contribute to these differences^77–80^.

Genes that are used repeatedly for evolutionary change have sometimes been described as “hotspots” of evolution^81^. Although hotspot loci have been reported in many species, the genomic, developmental, and ecological features that contribute to recurrent use of particular genes are still incompletely understood. In stickleback, many loci appear to be reused because of selection on ancient standing variants that already pre-exist at low frequency in marine populations^21,22^. The low frequency in marine populations is likely maintained by occasional hybridization between anadromous and resident freshwater fish, providing a continuing source of pre-existing variants that are available for adaptation of marine fish when they colonize freshwater habitats^82^. Similar reuse patterns may exist for many other loci underlying recurrent traits in stickleback^24–26,83^.

Genes may also be reused in evolution due to recurrent *de novo* mutations occurring independently at the same locus in multiple populations, and some researchers would restrict the term “hotspot” to such cases^81^. This pattern of repeated *de novo* mutations has previously been identified at the stickleback *PITX1* locus, where independent deletions of the same tissue-specific enhancer have led to recurrent evolution of pelvic skeletal reduction in multiple freshwater lakes^84^. Particular sequences within this enhancer have properties resembling fragile sites in human chromosomes^84^, and both elevated mutation rates and the strong phenotypic consequences of enhancer deletions likely explain why *PITX1* is used repeatedly when environmental conditions favor pelvic reduction^85^.

Our results suggest that the *MYH3C* cluster is an evolutionary hotspot in stickleback that is evolving through a combination of both ancient standing variants and more recent *de novo* mutations. The consistent SNP divergence between marine and freshwater *MYH3C* haplotypes, and the single microhomology breakpoint junction shared in multiple freshwater haplotypes, suggest that an ancestral 4-copy haplotype originally emerged through a single mutational event. Hastings et al.^57^ proposed such MMBIR events would be a driving force in evolution and disease by generating low copy repeats with the extended homology required for NAHR, leading to additional structural rearrangements in future generations. Indeed, we observed signatures of subsequent NAHR in 5-copy and 6-copy haplotypes as well as evidence for least three independent NAHR events in our sequenced freshwater myosin assemblies (two 4- to 5-copy and one 5- to 6-copy expansion). Thus, the initial 3- to 4-copy expansion in stickleback may have primed the *MYH3C* locus for future expansions and contractions. Rapid and repeated evolution of copy numbers in post-glacial freshwater stickleback likely occurs through a combination of selection on pre-existing myosin haplotypes, plus additional *de novo* changes occurring by recombination between expanded haplotypes. Such NAHR events are known to occur at frequencies 100 to 10,000 times higher than the rate of spontaneous mutations at single DNA base pairs^3^.

High intrinsic mutation rates may be a particularly important factor that biases reuse of hotspot loci during rapid evolutionary change in stickleback. Like many vertebrates, stickleback in postglacial lakes have evolved with smaller population sizes within a limited number of generations compared to typical microbial or invertebrate systems. These conditions probably favor evolution based either on pre-existing variants, or on *de novo* mutations that arise at high intrinsic rates at particular loci in the genome^85^. Many previous scans for recurrent adaptive loci in stickleback have searched for genomic windows that show shared nucleotide changes across multiple populations with similar phenotypes^23–26,86^. While such genomic patterns are straightforward to recognize, they are biased towards detecting recurrent evolution based on older variants that pre-exist in marine ancestors. CNVs provide an important additional source of high frequency *de novo* variation during adaptive evolution, and the current results should encourage more detailed studies of the structure and function of *MYH3C* alleles among populations, as well as searches for additional loci that evolve via recurrent copy number changes in particular environments^29^.

## RESOURCE AVAILABILITY

### Lead contact

Requests for further information and resources should be directed to and will be fulfilled by the lead contact, David Kingsley.

### Materials availability

Plasmids generated in this study have been deposited to Addgene (#250495, #250496).

### Data and code availability

- All raw sequencing data generated from this study have been deposited at the NCBI sequence read archive (SRA) as PRJNA1375864 and are publicly available as of the date of publication.
- *MYH3C* region assemblies generated from this study have been deposited at Genbank (accessions listed in Table S2) and are publicly available as of the date of publication.
- This paper analyzes existing, publicly available data, accessible at the SRA as: PRJNA247503, PRJNA671824, and additional accessions from the SRA and Genbank listed in Table S2.
- All original code is publicly available at Zenodo (doi: 10.5281/zenodo.18666765) as of the date of publication.
- Additional data files are publicly available at Zenodo (doi: 10.5281/zenodo.18666843) as of the date of publication.
- Any additional information required to analyze the data reported in this paper is available from the lead contact upon request.

## ACKNOWLEDGMENTS

We thank Aaron Daugherty, Julia Wucherpfennig, Gavin Sherlock, Anne Villeneuve, Andy Fire, Anne Brunet, Alex Urban, Matthew McCoy, and all members of the Kingsley lab for useful discussions; Ivan (Vanya) Zheludev and the Fire lab for assistance with early Nanopore sequencing experiments; Veronica Behrens and Katelyn (Katie) Sanko for assistance with BLAU sample dissections; and the Peichel lab for sharing BEPA BAC clones. Long-read sequencing data were collected at the University of Washington Long Reads Sequencing Center. This work was supported in part by a Stanford DARE predoctoral fellowship (R.R.D.), an American Cancer Society postdoctoral fellowship (E.E.H.), National Science Foundation Graduate Research Fellowships (A.M.Y., G.A.K., K.T.X), a Stanford CEHG Fellowship (H.I.C.), an Institutional National Research Service Award (T32) from the National Human Genome Research Institute (5T32HG000044-22, A.M.Y.), a Center of Excellence in Genomic Sciences grant from the National Institutes of Health (HG5P50HG002568, R.M.M. and D.M.K.), the Newcomb College Institute of Tulane University (D.C.H), National Science Foundation grants (DEB-0322818, DEB-0919184, M.A.B.), and a National Institutes of Health grant (1R01GM124330-0, M.A.B.). David Kingsley is an investigator of the Howard Hughes Medical Institute.

## AUTHOR CONTRIBUTIONS

A.M.Y., R.R.D., E.E.H., Y.F.C., F.C.J., and D.M.K. conceived the studies; E.H., R.R.D., and K.T.X carried out initial qPCR experiments and designed constructs; G.A.K. generated transgenic stickleback and phylogenetic trees of populations; T.R.H. generated the F2 cross and linkage map; H.Z. and A.A.P. carried out early myosin expression analyses; J.G., J.S. and R.M.M. sequenced and assembled the SALR myosin BAC clone; A.M.Y. and E.G.O. isolated and sequenced the BEPA myosin BAC clones and performed the allele-specific expression experiment. D.S., D.C.H., T.E.R., B.J. and M.A.B. provided key samples and suggestions based on extant and recently evolving populations; A.M.Y., R.R.D., H.I.C., M.A.B. and B.J. collected Alaskan and Icelandic samples; A.M.Y., E.G.O., and S.D.B. prepared and managed DNA samples; C.L. and E.A. provided RABS marine genome information; A.M.Y and R.R.D. carried out most experimental studies and analyses; M.L.R., J.A.B., and R.R.D. investigated swimming behavior; and A.M.Y, R.R.D. and D.M.K wrote the paper with input from all coauthors.

## STAR★METHODS

### KEY RESOURCES TABLE

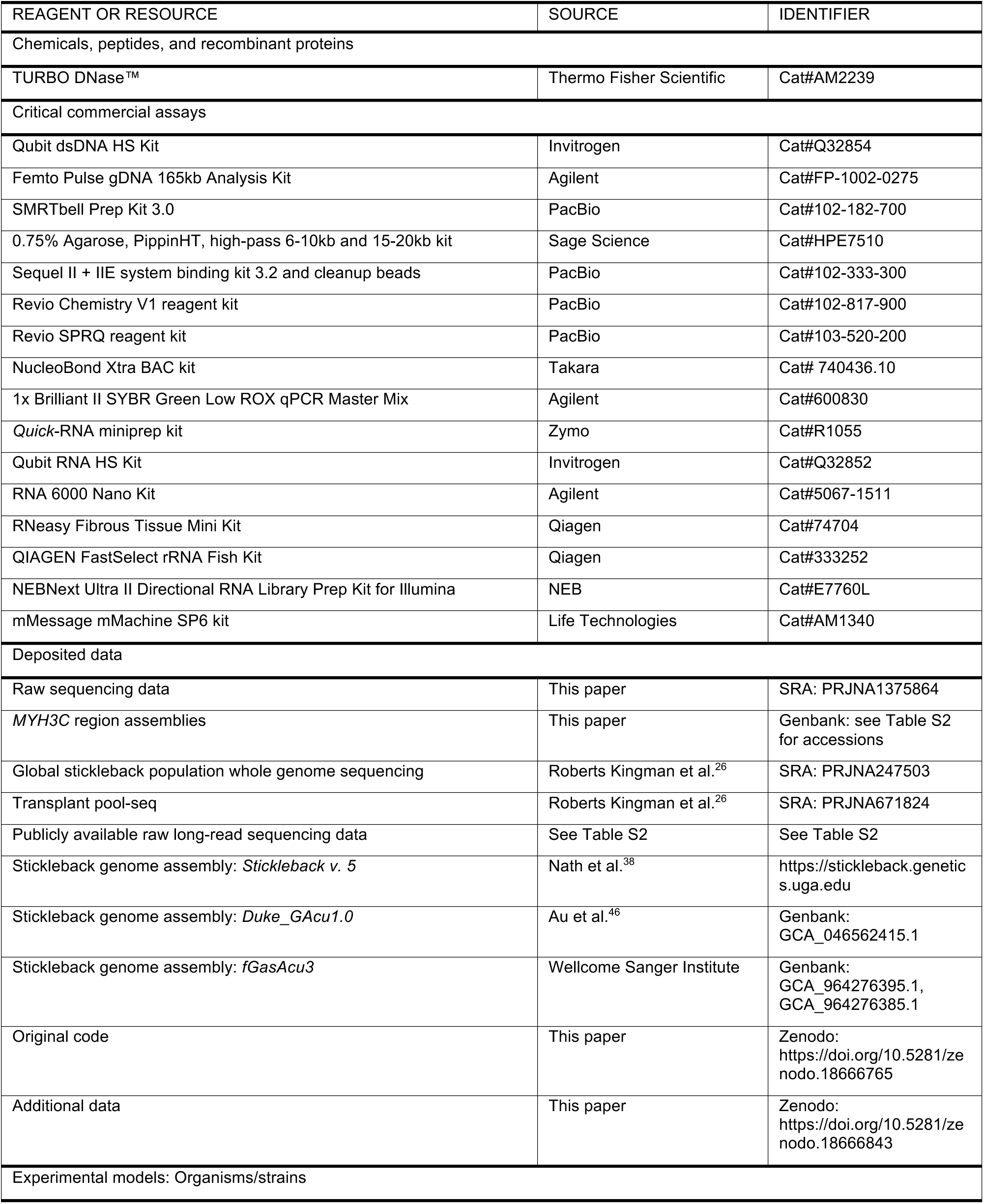
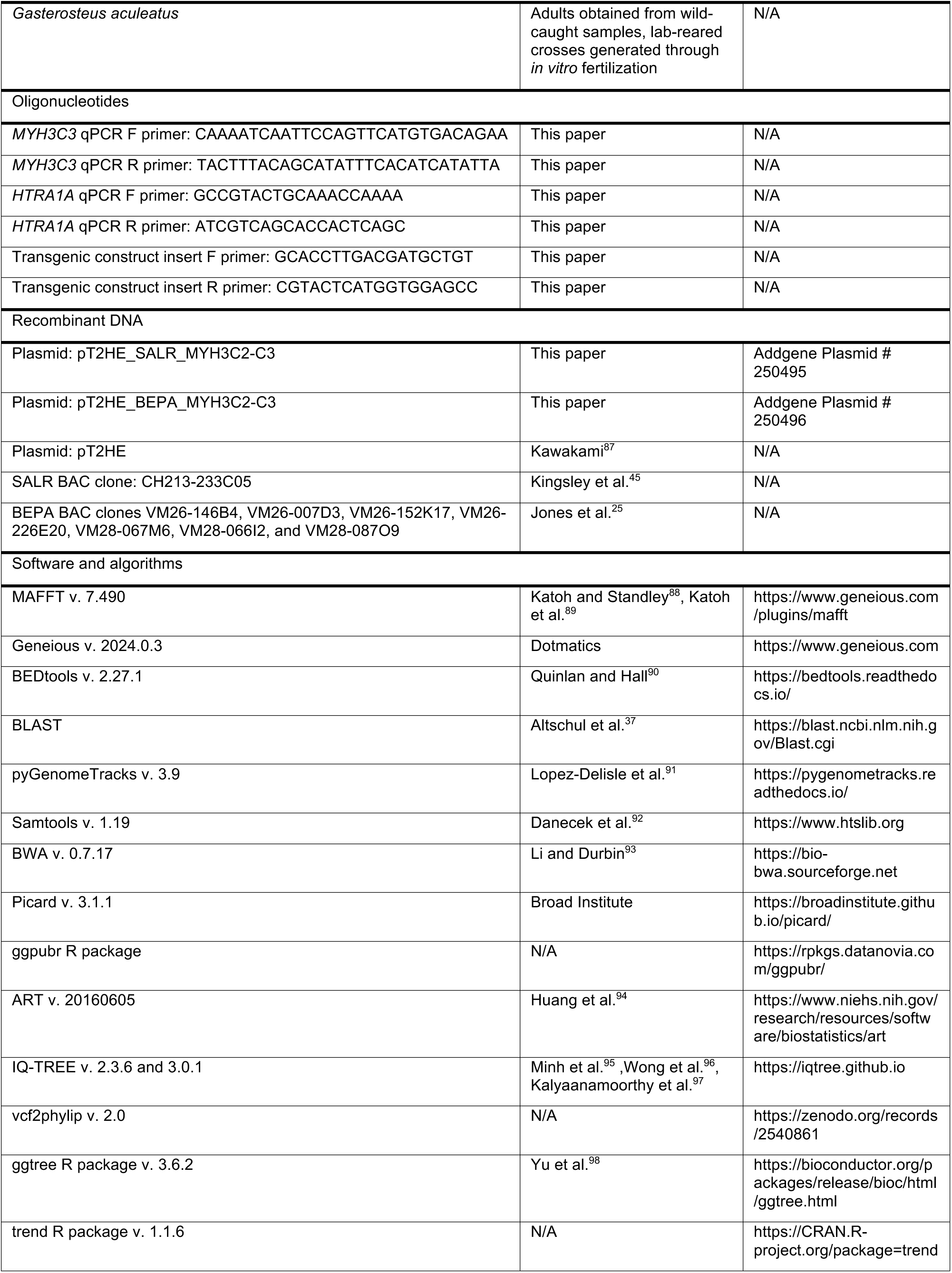
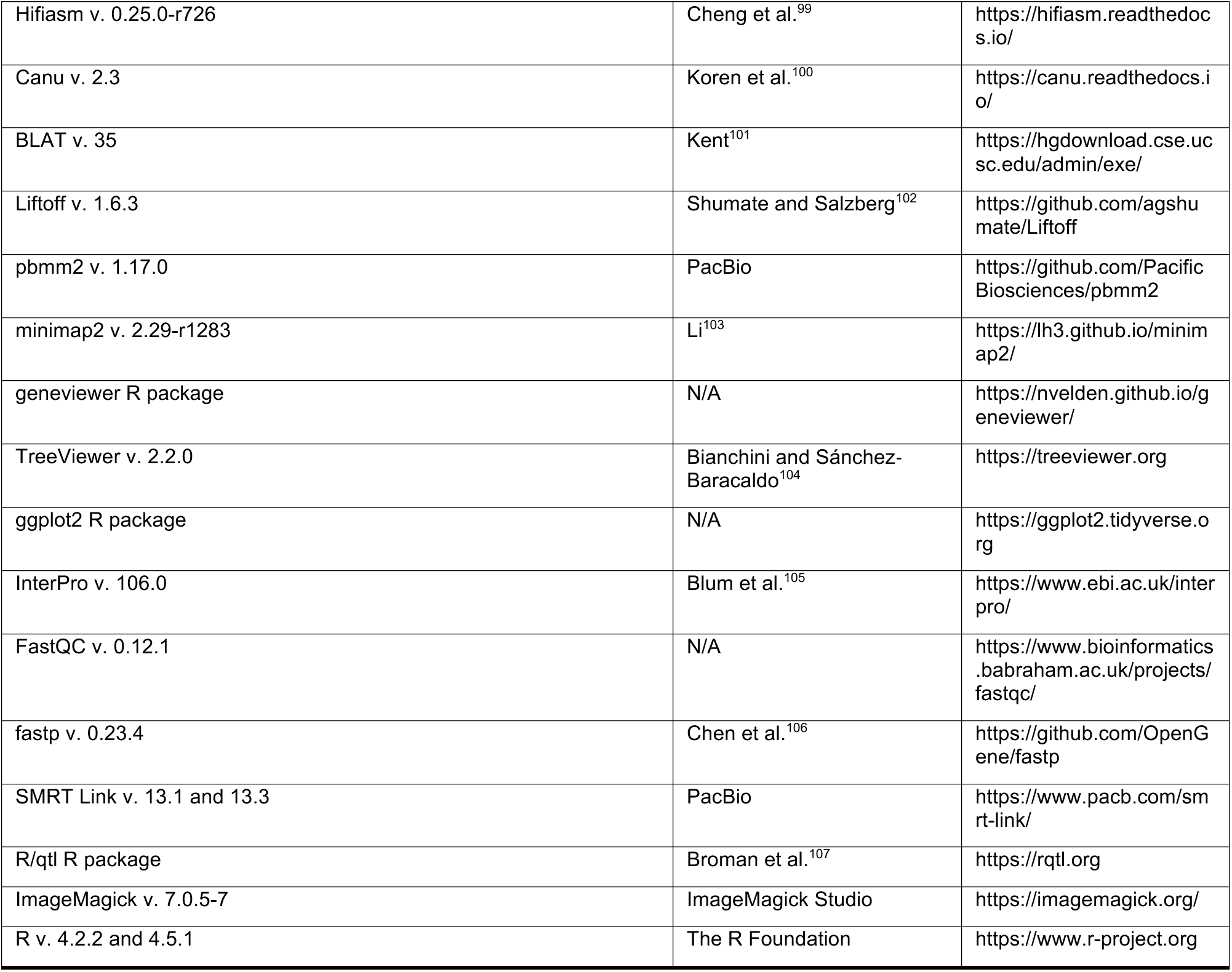

### EXPERIMENTAL MODEL AND STUDY PARTICIPANT DETAILS

#### Ethical compliance and animal care

All stickleback care was performed in accordance with the Guide for the Care and Use of Laboratory Animals of the National Institutes of Health, following protocols approved by the Institutional Animal Care and Use Committee of Stanford University (protocol #13834). Wild stickleback from Alaska and Iceland were collected in unbaited minnow traps and euthanized in MS-222 (see Table S2 for collection locations).

#### Allele-specific expression experiment

Two lab-reared marine RABS females (siblings) were crossed with one lab-reared BEPA male by *in vitro* fertilization to produce two clutches of RABS x BEPA F1 hybrids. Fish were raised under standard lab conditions at 18°C. At 27 dpf, approximately half the fish from each clutch were randomly selected for an early developmental time collection. Of these, fish were randomly split into two groups, one that continued under the same 18°C temperature condition, and another that was acclimated to 3°C. This group was acclimated to 8°C overnight, then transferred into net breeders in a tank with a chiller that was lowered by 2°C every 1.5 hours until the final water temperature reached 3°C. At 31 dpf, fish from both 18°C and 3°C conditions were euthanized in MS-222. A caudal fin clip was stored in 70% ethanol and used for IDH sex genotyping^49^ while the remainder of the body was flash-frozen in liquid nitrogen and stored at 80°C. The remaining fish were raised at 18°C until they were 5 months old (152 dpf) when half of the fish were acclimated to 8°C overnight. These fish were then transferred into a tank with a chiller and the temperature was lowered by 1°C per day until reaching a final temperature of 3°C. At 161 dpf, all fish were euthanized in MS-222. Pectoral adductor muscle, pectoral abductor muscle (including attached skin and cartilage due to small size), and flank muscle were collected, flash frozen in liquid nitrogen, and stored at −80°C. A caudal fin clip was collected in 70% ethanol and used for IDH sex genotyping^49^.

### METHOD DETAILS

#### Custom annotation of the myosin cluster

The *stickleback v. 5* reference genome^38^ and annotations were modified and used for all analyses unless otherwise noted. *MYH3C* annotations were manually corrected from NCBI and Ensembl annotations using visual inspection of mapped Kinnex reads from allele-specific experiment samples, one male fish from Cheney Lake, and one male stickleback from BLAU (see below). Myosin CNV breakpoints were identified by self-alignment using MAFFT v. 7.490^88,89^ with default parameters and dot plot analysis implemented in Geneious v. 2024.0.3 (https://www.geneious.com). For read depth analysis, duplication region 2 (ChrXIX:2683412-2701087; Figure 1A) was hardmasked using BEDtools v. 2.27.1^90^ and ChrY was removed to avoid multimapping reads and permit pileup of reads from duplicated copies at a single locus. Global sensitive EcoPeak and Cheney-Scout-Loberg Sensitive TempoPeak coordinates described by Roberts Kingman et al.^26^ were transferred from *gasAcu1-4* to *stickleback v. 5* using LiftOver (UCSC tools v. 469)^108^. NCBI annotated gene LOC120809426 was identified as *HARBI1* based on BLASTP search against the non-redundant protein sequences database (nr)^37^. The *MYH3C* genome region was plotted using pyGenomeTracks v. 3.9^91^.

#### Ecotypic read depth analysis

*C3* copy number changes were detected by *C3* read depth analysis of whole genome sequence data from globally-distributed stickleback populations (SRA accession PRJNA247503)^26^. Reads were extracted from *gasAcu1-4*-aligned bam files and separated into read groups using Samtools v. 1.19^92^. Reads were realigned to the modified *stickleback v. 5* reference using bwa v. 0.7.17^93^ and sorted using Samtools v. 1.19^92^. Picard v. 3.1.1^109^ was used to mark duplicates and annotate read groups. Aligned bams from read groups were merged into one file per individual and duplicates were marked again using Picard v.3.1.1^109^. Read depth was calculated using Samtools coverage with a minimum MAPQ=3 and requiring primary alignments^92^. *C3* normalized read depth was calculated by dividing *C3* read depth by the mean autosomal read depth. Eight samples were identified as potential males based on a ChrXIX read depth approximately 25% lower than the autosomal chromosomes and were removed from further analysis. Samples originally described by Jones et al. were also excluded due to shorter read lengths which affected mappability and depth^25^. Remaining samples included in the Pacific (c150), Northern Europe (c151), and California freshwater with Pacific marine (c153) sample subsets designated by Roberts Kingman et al.^26^ were used for plotting and statistical comparison of freshwater vs. marine read depth. Marine vs. freshwater and hybrid zone *C3* copy differences were evaluated using a Wilcoxon rank sum test and latitude correlations were determined using Pearson correlation implemented by the ggpubr R package^110^. Simulated reads were used to interpret allele copy number from *C3* read depth. We replaced the myosin region (including *NLCR5* through *SYT19*) in either the freshwater *stickleback v. 5* (without chrY) reference or the marine *Duke*_*GAcu1.0* reference^46^ with the equivalent regions from 3- to 6-copy myosin assemblies (see below). Paired-end 76 bp reads based on the HiSeq2000 with a mean fragment size of 150 bp and standard deviation of 50 bp at 8x depth were simulated using ART v. 20160605^94^ and read depth analysis was performed as described above. When simulated reads from multiple assemblies for a CNV allele were available, final *C3* read depth was averaged before plotting.

#### Neutral phylogenetic tree construction

Neutral genomic regions were defined as parts of the genome located at least 5 Mb from any sensitive EcoPeak (Pacific or global) identified by Roberts Kingman et al^26^. The variant call file (.vcf) from Roberts Kingman et al.^26^ was filtered to include only samples from Figure 1B (this paper) and to exclude all variants not falling within this conservative definition of neutral genomic regions. Filtered variants were then randomly subsampled to 100,000 variants (with minor allele frequency > 0.02) to ensure reasonable tree building runtime. The variant file was converted from VCF to PHYLIP format^111^ and IQ-TREE3^95,96^ was used to generate a log-likelihood maximized tree of genetic relationships between the neutral genomic regions of these samples. The resulting phylogenetic tree was visualized using the ggtree R package^98^.

#### Transplant pool-seq analysis

Marine threespine stickleback that had entered freshwater to breed in Rabbit Slough, AK were transplanted into freshwater lakes (Cheney Lake, Scout Lake) and a third lake (Loberg Lake) was naturally colonized by Rabbit Slough or closely related marine fish^44^. The three lakes were sampled annually, and sequenced in pools of 39-200 fish as previously described^26,43^. Pool-seq data (SRA accession PRJNA671824) were realigned to the modified *stickleback v.5* reference and read depth was calculated as described above for ecotypic depth analysis with the following differences: (1) ChrY was included in the reference, and (2) normalized *C3* read depth was calculated by dividing *C3* read depth by the mean ChrXIX read depth to normalize for different sex ratios in each set of pooled fish. A one-sided Mann-Kendall trend test (alternative hypothesis: true S > 0; trend R package v. 1.1.6^112^) was applied to each population to evaluate increasing *C3* read depth over time, the prediction based on previously known read depth patterns among marine and freshwater populations. Allele frequencies of the most significant SNP within the myosin region Sensitive TempoPeak, ChrXIX:2,744,769 (*gasAcu1-4* reference), were previously determined by Roberts Kingman et al.^26^ and plotted.

#### High molecular weight (HMW) DNA extractions

To collect blood for high molecular weight DNA extraction, fish were anesthetized with MS-222, the caudal peduncle was cut, and the fish was held vertically with the posterior end just submerged in 1.5 ml of 0.85x SSC buffer on ice. Blood samples were stored at 4°C for up to one week and divided equally into two aliquots using wide-orifice pipets. Each aliquot was centrifuged at 2000xg for 2 min at 4°C and the supernatant was removed. Blood pellets were frozen on dry ice and stored at −80°C. For BLAU, blood was instead collected directly from the cut caudal peduncle onto glass slides, air dried overnight, then stored at room temperature in coin envelopes in bags with desiccant.

Blood samples (either thawed blood pellets or dried blood scraped from slides) were digested in 10 μl of Proteinase K and 600 μl of lysis buffer (10 mM Tris, pH 8, 100 mM NaCl, 10 mM EDTA, 0.5% SDS) and incubated for 1-2 hours at 55°C. The cell lysate was combined with 600 μl of phenol:cholorform:isoamyl alcohol 25:24:1, gently mixed by rotating for 1 hour, and centrifuged for 2 min at 3000xg. The aqueous layer was transferred to a new tube and the phenol:chloroform step was repeated if needed until the aqueous portion was colorless. The aqueous layer was mixed with 600 μl of chloroform, rotated for 5 min, then centrifuged for 2 min at 3000xg, and repeated for two total chloroform washes. The aqueous layer was mixed with 1 mL of −20°C 100% ethanol and 30 μl of 3M NaOAc (pH 5.2), rotated for 10 min until a ball of precipitated DNA was visible, then centrifuged for 1 min at 13,500xg. The supernatant was removed and replaced with 500 μl of 70% ethanol. Each sample was centrifuged for 1 min at 13,500xg, the supernatant was removed, and the DNA pellet was air dried for 10 min. Two hundred μl of Qiagen EB buffer was added to the DNA pellet and left overnight at 4°C to resuspend. All phenol chloroform and chloroform centrifugation steps were performed in microcentrifuge tubes with Corning vacuum grease which separated the aqueous and organic phases and permitted pouring of the aqueous layer into new tubes. All other handling of HMW DNA was performed with wide-orifice pipette tips.

#### PacBio HiFi sequencing

PacBio HiFi data were generated per manufacturer’s recommendations at the University of Washington Long Reads Sequencing Center. At all steps, quantification was performed with Qubit dsDNA HS (Invitrogen, Q32854) measured on DS-11 FX (Denovix) and size distribution checked using FEMTO Pulse (Agilent, M5330AA & FP-1002-0275). HMW DNA was sheared with Megaruptor 3 (Diagenode, B06010003 & E07010003) using settings 28/31 or 28/30 (depending on original length distribution) and used to generate PacBio HiFi libraries via the SMRTbell Prep Kit 3.0 (PacBio, 102-182-700) using barcoded adapters (PacBio, 102-009-200). Size selection was performed with Pippin HT using a high-pass cutoff of 10-17 kbp (Sage Science, HTP0001 & HPE7510). BLAU was sequenced on the Sequel II platform on SMRT Cells 8M (PacBio, 101-389-001) using Sequel II Sequencing Chemistry 3.2 (PacBio,102-333-300) with 2-hour pre-extension and 30-hour movies. BEPA, CMCB, DNSE, ECHO, JADE, and KFSY were sequenced on the Revio platform on SMRT Cells 25M with Revio Chemistry V1 (PacBio, 102-817-900) with Adaptive Loading and 30-hour movies.

#### Genome assembly and annotation

*De novo* genome assembly was performed on PacBio HiFi CCS reads from BEPA, CMCB, DNSE, ECHO, JADE, KFSY, BLAU (see above) and BOULxBDGB (SRA accession SRR16093180) using Hifiasm v. 0.25.0-r726^99^. Other publicly available stickleback whole genome long read datasets based on PacBio CLR reads or Oxford Nanopore sequencing were downloaded and assembled using Canu v. 2.3^100^. Publicly available stickleback assemblies derived at least partially from long read sequencing methods were also analyzed (Table S2). For each assembly, contigs containing *MYH3C* copies were identified using BLAT v. 35^101^. NCBI and custom *MYH3C* annotations were transferred from *stickleback v. 5* to each assembly using Liftoff v. 1.6.3 allowing extra copies with 98% minimum sequence identity to be annotated^102^. Reads were remapped to their corresponding assemblies using either pbmm2 v. 1.17.0 (https://github.com/PacificBiosciences/pbmm2; accessed on 3 June 2025) for PacBio reads or minimap2 v. 2.29-r1283^103^ for Oxford Nanopore reads and visually inspected for quality. Any assemblies that showed uneven read coverage or high frequency sequence differences in reads relative to the assembly in the *MYH3C* region were removed from further analysis. Copy number haplotypes were visualized using the geneviewer R package^113^.

#### SALR BAC sequencing

Paired-end sequence reads were used to identify a *MYH3C* region clone from a previously described Bacterial Artificial Chromosome (BAC) library made from the SALR marine population^25,45^. The CH213-233C05 clone was sequenced to completion as described^49^.

#### BEPA BAC sequencing

Paired-end sequence reads were used to identify seven BAC clones fully spanning the *C3* copies from two BAC libraries made from the BEPA stickleback reference genome fish^25^. Clones VM26-146B4, VM26-007D3, VM26-152K17, VM26-226E20, VM28-067M6, VM28-066I2, and VM28-087O9 were purified using the NucleoBond Xtra BAC kit (Takara, 740436.10). Whole plasmid sequencing was performed by Plasmidsaurus using Oxford Nanopore Technology with custom analysis and annotation. Individual reads were annotated with *MYH3C* gene copies using Live Annotate & Predict (85% similarity) in Geneious v. 2024.0.3 (https://www.geneious.com).

#### QTL mapping of C3 DNA copy number

A wild-caught freshwater female (Boulton Lake, BC, Canada) and marine male (Bodega Bay, CA) stickleback were crossed *in vitro* to produce F1s, which were then intercrossed to produce multiple F2 families. Four-hundred forty fish from the largest F2 family were genotyped, and phasing and linkage map construction were performed as described by Wucherpfennig et al.^47^ In this analysis, fish with fewer than 600 genotype calls and markers that had genotype calls in fewer than 320 samples were excluded. Non-informative markers and makers showing irregularities in segregation ratios or inconsistent placement on the linkage map were removed, and the linkage map was rebuilt using a reduced set of individuals and markers (341 F2 fish, 448 markers).

To assess DNA copy number, primers were designed to amplify a 164 bp product specific to *C3* (5’-CAAAATCAATTCCAGTTCATGTGACAGAA-3’ and 5’-TACTTTACAGCATATTTCACATCATATTA-3’). *HTRA1A*(5’-GCCGTACTGCAAACCAAAA-3’ and 5’-ATCGTCAGCACCACTCAGC-3’) was used as an intra-sample comparison gene and showed no apparent copy number variation in the depth analysis of global stickleback populations described above. PCR reactions were performed in 25 μl reactions, with 1x Brilliant II SYBR Green Low ROX qPCR Master Mix (Agilent, 600830), 0.4 μM of each of forward and reverse primer, and 5 ng of DNA. All qPCR was performed using a QuantStudio 5 Real-Time PCR System (Thermo Fisher) with the following protocol: (1) one cycle of 15 minutes at 95 °C (2) 40 cycles of 10 seconds at 95 °C and 30 seconds at 60°C. Analysis of cycle thresholds (Ct) used the relative comparative Ct method (2^−ΔΔCt^)^114^. Means for each sample were calculated from three independent replicates for each individual in the study. A standard fish sample from BEPA with 6 diploid copies of *C3*, was used to calibrate every qPCR plate and estimate the absolute number of *C3* copies for samples.

*C3* DNA copy number was compared to 214 mapped genomic SNP markers that were homozygous for alternative alleles in the grandparents for QTL analysis in R/qtl^107^. Significant chromosomal loci were identified via a single-QTL model with a threshold based on 10,000 ‘scanone’ permutations. Haley-Knott regression was used as the interval mapping method, and a normal model was used for the copy number phenotype.

#### Phylogenetic analysis of MYH3C orthologs

Putative *MYH3C* orthologs were identified using BLASTP searches and Ensembl release 110-listed orthologs of stickleback *MYH3C* annotations (ENSGACG00000002902, ENSGACG00000002933, ENSGACG00000002955, ENSGACG00000003003). The nucleotide sequences from the protein-coding regions of the canonical transcripts from *Danio rerio* (*myhz1.1*, ENSDARG00000067990; *myhz1.2*, ENSDARG00000067995; *myhz1.3*, ENSDARG00000067997; *myhz2*, ENSDARG00000012944; *myhc4*, ENSDARG00000035438; *myha*, ENSDARG00000095930), *Orzias melastigma* (*C1*, ENSOMEG00000014116; *C2*, ENSOMEG00000014025; *C3*, ENSOMEG00000013940), and *Pungitius pungitius* (*C1*, ENSPPNG00035026282; *C2*, ENSPPNG00035026670; *C3*, ENSPPNG00035027195) putative *MYH3C* orthologs were aligned using MAFFT v. 7.490^88,89^ with default parameters implemented in Geneious v. 2024.0.3 (https://www.geneious.com). Maximum likelihood analysis was performed using IQ-TREE v. 2.3.6 with default parameters and 100 non-parametric bootstraps^95,97^. The resulting phylogenetic tree was visualized using TreeViewer v. 2.2.0^104^.

#### Protein sequence analysis

Protein sequences for all stickleback *MYH3C* copies were aligned using MAFFT v. 7.490^88,89^ with default parameters implemented in Geneious v. 2024.0.3 (https://www.geneious.com). Divergent amino acid positions were extracted from the alignment using custom scripts and plotted in R using the ggplot2 R package^115^. Positions of protein domains were determined by searching the BEPA C3A protein sequence in InterPro v. 106.0^105^. PROSITE annotations for the SH3 domain (33-83 aa; PS51844), myosin motor domain (87-774 aa; PS51456), and IQ motif (777-806 aa; PS50096) and the Pfam annotation for the myosin tail (842-1919 aa; PF01576) were plotted.

#### NAHR breakpoint analysis

The full 17.6 kb duplicated regions from each 5-copy and 6-copy assembly were aligned using MAFFT v. 7.490^88,89^ with default parameters implemented in Geneious v. 2024.0.3 (https://www.geneious.com). Regions with gaps were removed from the alignment and divergent positions were identified using custom scripts. Within each assembly, only positions that differed between the duplication regions spanning *C3A* and *C3L* were plotted. All nucleotide positions in intermediate duplication regions (B, C) were then identified as matching the A or L sequence. Regions were designated as candidate crossover regions where the broad pattern of variants present in duplication regions B or C transitioned from matching the L sequence to matching the A sequence.

#### Cheney and BLAU RNA samples

To annotate ChrY *MYH3C* copies, abductor pectoral muscle was dissected from one wild-caught male adult stickleback each from Cheney Lake and BLAU. Tissue samples were stored in RNAlater at 4°C for up to 3 weeks, then stored at −80°C. RNA was extracted using the *Quick*-RNA miniprep kit (Zymo, R1055). Tissue samples were homogenized in 600 μl of RNA lysis buffer using MP Fastprep 2x30s program with one ¼” ceramic bead and a 5 min rest between. RNA was eluted in 50 μl RNase-free water. RNA concentration was determined using the Qubit RNA HS kit (Invitrogen, Q32852) and integrity was checked by Bioanalyzer using the RNA 6000 Nano Kit (Agilent, 5067-1511).

#### RNA-seq

Six RABS x BEPA F1 hybrid females were randomly selected for each condition (3°C vs. 18°C temperature and 1 month vs. 5 month developmental stage), except the 3°C, 5 month condition where only 5 females were available (see Section: Allele-specific expression experiment). RNA was extracted using the RNeasy Fibrous Tissue Mini Kit (Qiagen, 74704). Tissue samples were homogenized in 300-900 μl of Buffer RLT (depending on tissue size) using the MP Fastprep 2x30s program with one ¼” ceramic bead and a 5 min rest between. RNA was eluted in 30 μl RNase-free water.

RNA sample QC, library preparations, sequencing reactions, and initial bioinformatic analysis were conducted at GENEWIZ, LLC./Azenta US, Inc (South Plainfield, NJ, USA). Total RNA samples were quantified using the Qubit 4.0 Fluorometer (Invitrogen, Q33226) and RNA integrity was checked with 4200 TapeStation (Agilent Technologies, G2991BA). Samples were first treated with TURBO DNase (Thermo Fisher Scientific, AM2239) to remove DNA contaminants. Then rRNA depletion was performed using the QIAGEN FastSelect rRNA Fish Kit (Qiagen, 333252) according to the manufacturer’s protocol. A strand-specific RNA sequencing library was prepared by using the NEBNext Ultra II Directional RNA Library Prep Kit for Illumina following the manufacturer’s instructions (NEB, E7760L). Briefly, the enriched RNAs were fragmented for 8 minutes at 94°C. First strand and second strand cDNA were subsequently synthesized. The second strand of cDNA was marked by incorporating dUTP during the synthesis. cDNA fragments were adenylated at 3’ends, and indexed adapter was ligated to cDNA fragments. Limited cycle PCR was used for library enrichment. The incorporated dUTP in the second strand cDNA quenched the amplification of the second strand, which helped to preserve the strand specificity. The sequencing library was validated on the Agilent TapeStation (Agilent Technologies, G2991BA) and quantified by using Qubit 4.0 Fluorometer (Invitrogen, Q33226) as well as by quantitative PCR (KAPA Biosystems). The sequencing libraries were multiplexed and clustered on the flowcell. After clustering, the flowcell was loaded on the Illumina NovaSeq instrument according to manufacturer’s instructions. The samples were sequenced using a 2x150 Pair-End (PE) configuration. The NovaSeq Control Software conducted image analysis and base calling. Raw sequence data (.bcl files) generated by the sequencer were converted into fastq files and de-multiplexed using Illumina’s bcl2fastq 2.20 software. One mismatch was allowed for index sequence identification.

Raw reads were checked for quality using FastQC v. 0.12.1^116^, then trimmed (fastp v. 0.23.4^106^) to remove any residual adaptors and the first 13 bases of each read which showed nucleotide composition bias from random hexamer priming. Because of the high sequence identity between *MYH3C* transcripts, traditional reference mapping approaches of RNA-seq reads result in mismapping and reference biases for quantifying *MYH3C* expression. We instead identified two types of 27-bp *k*-mers designed to specifically target known *MYH3C*-distinguishing variants within exons based on alignment of RABS and BEPA genomic assemblies. First, *MYH3C* copy expression (MCE) 27-mers target the same region in all *C1*, *C2*, and *C3* copies, but contain variants that distinguish *C1* from *C2* from *C3*, and no variants that distinguish the BEPA and RABS allele. Because these *k*-mers detect equivalent regions in all *MYH3C* copies and account for technical differences in read coverage along the transcript, they were used for comparison of the relative total expression of *C1*, *C2*, and *C3* across conditions. To distinguish expression from the BEPA or RABS alleles, we designed allele-specific expression (ASE) *k*-mers which contain at least one variant that distinguishes a myosin copy from the others (i.e. *C3* distinguished from *C1* and *C2*) as well as a variant that distinguishes the BEPA allele from the RABS allele. To check *k*-mer quality, we BLAT-searched (BLAT v. 35^101^) the *k*-mers against the RABS and BEPA reference genomes and removed any *k*-mers with either a perfect off-target match or a substantial number (>5) of off-target matches with 1-bp difference. *K*-mers were also compared to mapped Kinnex reads (see below) to confirm the expected distinguishing variants were present in the F1 hybrids and no additional variants that would prevent *k*-mer detection were present. We also required that *k*-mers be at least 100 bp from the end of a transcript as read coverage was much lower at transcript ends. This resulted in seven MCE, eight *C1* ASE, four *C2* ASE, and nine *C3* ASE sets of 27-mers (Table S4). Raw reads were searched for exact matches to each 27-mer and a read pair was counted only once if there was a match in the forward, reverse, or both reads. Reads per million (RPM) was calculated (*k*-mer read detections / total read pairs * 1,000,000) for MCE 27-mers. To calculate ASE, we required that a sample have at least one BEPA and RABS read each, a minimum of 100 total reads, and all *k*-mers for that *MYH3C* copy detected. The proportion of the BEPA allele was calculated by dividing the BEPA 27-mer counts by the sum of the BEPA and RABS counts, then averaging across all 27-mers for each *MYH3C* copy. For each tissue, ASE was evaluated with a one-sample Wilcoxon rank sum test (mu = 0.5) and temperature ASE was tested with a two-sample Wilcoxon rank sum test implemented in the ggpubr R package^110^.

#### Kinnex

The Cheney and BLAU pectoral muscle RNA samples and two F1 hybrid RNA samples for each developmental stage, tissue, and temperature condition were used to prepare Kinnex libraries. Total RNA was quality checked using UV-Vis spectroscopy (Denovix DS-11 FX) and Agilent Bioanalyzer 2100 using the Total RNA Nano 6000 kit (Agilent, G2939A & 5067-1511). Kinnex full-length RNA libraries were generated per manufacturer’s recommendations (PacBio, 103-072-000). Samples were sequenced on the Revio platform on SMRT Cells 25M with Revio Chemistry V1 (PacBio, 102-817-900) for Cheney and BLAU samples, or Revio SPRQ (PacBio, 103-520-200) for F1 hybrid samples, with Adaptive Loading and 30-hour movies. Data were postprocessed using SMRT Link v13.1 or 13.3 with the “Read Segmentation and Iso-Seq” pipeline to segment and classify reads. Full-length non-chimeric (FLNC) reads from Cheney and BLAU samples were mapped to the modified *stickleback v.5* reference using pbmm2 v. 1.17.0 (https://github.com/PacificBiosciences/pbmm2; accessed on 3 June 2025) and used to adjust *MYH3C* annotations. To facilitate accurate mapping of *MYH3C* reads from F1 hybrids, we created a composite marine and freshwater reference comprised of the marine RABS *Duke_GAcu1.0* reference^46^ and the freshwater *stickleback v. 5* reference (no ChrY) with the 5-copy BEPA haplotype replacing the *MYH3C* region. F1 hybrid FLNC reads were aligned to the composite reference genome using pbmm2 v. 1.17.0 (https://github.com/PacificBiosciences/pbmm2; accessed on 3 June 2025) with a maximum intron length of 10,000 bp. Primary alignment reads fully spanning exons 3-35 (containing all 9 variants that can be used to distinguish BEPA *C3* copies) were extracted and counted to determine ratios of *MYH3C* copy expression.

#### Transgenic enhancer assays

An approximately 2.6-2.8 kb region spanning from 3’ end of *C2* to the 5’ end of *C3*, including the intergenic region was PCR amplified (5’-GCACCTTGACGATGCTGT-3’ and 5’-CGTACTCATGGTGGAGCC-3’) from the marine SALR BAC clone and the BEPA stickleback freshwater reference individual. These amplicons were then separately cloned into the pT2HE vector at the PspOMI site, upstream of an eGFP reporter gene (modified from Kawakami^87^). The resulting constructs, or an empty vector control, were microinjected along with mature *Tol2* transposase mRNA into fertilized stickleback eggs to generate transgenic fish as previously described^84^. The mature *Tol2* transposase mRNA was produced by *in vitro* transcription using the mMessage mMachine SP6 kit (Life Technologies, AM1340). Transgenic fry were imaged at 7.5 mm standard length, approximately 23 dpf. The pT2HE vector constitutively induces expression in the eye, providing a control site to assess successful injection and degree of transgene expression, which can vary in founder larvae due to different integration sites and degree of mosaicism. Images were stripped of all metadata, pooled, randomized, and distributed to three independent reviewers, who each assigned a score to individual fish from 1 (low) to 5 (high) for strength of fluorescence in the left eye, right eye, left flank muscle, and right flank muscle. Images were then unmasked, and normalized flank intensity scores (average flank divided by average eye, averaged across all viewers and both sides of the fish) were tabulated and compared by Wilcoxon rank sum test. Scores from all three reviewers were similar, with no outliers in any category of statistical analysis. Brightness and contrast adjustments were equally applied to representative images shown in Figure 4B using ImageMagick v. 7.0.5-7 (-brightness-contrast 25x35)^117^.

### QUANTIFICATION AND STATISTICAL ANALYSIS

Data were analyzed using R v. 4.2.2 or 4.5.1. The number of individuals is represented by *n*, except for Figure 2C, where *n* refers to the number of protein copies originating from individuals and multiple copies present in a single individual. Statistical tests used included a two-sided Wilcoxon rank sum test, Pearson correlation, a one-sided Mann-Kendall trend test, a hypergeometric test, a one sample Wilcoxon rank sum test, and a LOD score. Boxplots display the median (line), 25^th^-75^th^ percentiles (box), and 1.5 x IQR (whiskers).

## SUPPLEMENTAL INFORMATION

Document S1. Figures S1–S6, Tables S3 and S4, and supplemental references

Table S1. Global stickleback samples used for *C3* read depth calculations, related to Figures 1, S1, and S2

Table S2. Samples used for generating *MYH3C* assemblies, related to Figure 2

## REFERENCES

1. Conrad, D.F., Pinto, D., Redon, R., Feuk, L., Gokcumen, O., Zhang, Y., Aerts, J., Andrews, T.D., Barnes, C., Campbell, P., et al. (2010). Origins and functional impact of copy number variation in the human genome. Nature 464, 704–712. 10.1038/nature08516.

2. Redon, R., Ishikawa, S., Fitch, K.R., Feuk, L., Perry, G.H., Andrews, T.D., Fiegler, H., Shapero, M.H., Carson, A.R., Chen, W., et al. (2006). Global variation in copy number in the human genome. Nature 444, 444–454. 10.1038/nature05329.

3. Zhang, F., Gu, W., Hurles, M.E., and Lupski, J.R. (2015). Copy number variation in human health, disease and evolution. Annu. Rev. Genomics Hum. Genet. 10, 451–481. 10.1146/annurev.genom.9.081307.164217.

4. Nathans, J., Piantanida, T.P., Eddy, R.L., Shows, T.B., and Hogness, D.S. (1986). Molecular genetics of inherited variation in human color vision. Science 232, 203–210. 10.1126/science.3485310.

5. Lupski, J.R., Wise, C.A., Kuwano, A., Pentao, L., Parke, J.T., Glaze, D.G., Ledbetter, D.H., Greenberg, F., and Patel, P.I. (1992). Gene dosage is a mechanism for Charcot-Marie-Tooth disease type 1A. Nat. Genet. 1, 29–33. 10.1038/ng0492-29.

6. MacIntyre, D.J., Blackwood, D.H.R., Porteous, D.J., Pickard, B.S., and Muir, W.J. (2003). Chromosomal abnormalities and mental illness. Mol. Psychiatry 8, 275–287. 10.1038/sj.mp.4001232.

7. Sebat, J., Lakshmi, B., Malhotra, D., Troge, J., Lese-Martin, C., Walsh, T., Yamrom, B., Yoon, S., Krasnitz, A., Kendall, J., et al. (2007). Strong association of de novo copy number mutations with autism. Science 316, 445–449. 10.1126/science.1138659.

8. McCarroll, S.A., Huett, A., Kuballa, P., Chilewski, S.D., Landry, A., Goyette, P., Zody, M.C., Hall, J.L., Brant, S.R., Cho, J.H., et al. (2008). Deletion polymorphism upstream of *IRGM* associated with altered *IRGM* expression and Crohn’s disease. Nat. Genet. 40, 1107–1112. 10.1038/ng.215.

9. Perry, G.H., Dominy, N.J., Claw, K.G., Lee, A.S., Fiegler, H., Redon, R., Werner, J., Villanea, F.A., Mountain, J.L., Misra, R., et al. (2007). Diet and the evolution of human amylase gene copy number variation. Nat. Genet. 39, 1256–1260. 10.1038/ng2123.

10. Axelsson, E., Ratnakumar, A., Arendt, M.-L., Maqbool, K., Webster, M.T., Perloski, M., Liberg, O., Arnemo, J.M., Hedhammar, Å., and Lindblad-Toh, K. (2013). The genomic signature of dog domestication reveals adaptation to a starch-rich diet. Nature 495, 360–364. 10.1038/nature11837.

11. Rinker, D.C., Specian, N.K., Zhao, S., and Gibbons, J.G. (2019). Polar bear evolution is marked by rapid changes in gene copy number in response to dietary shift. Proc. Natl. Acad. Sci. U.S.A. 116, 13446–13451. 10.1073/pnas.1901093116.

12. Ishikawa, A., Kabeya, N., Ikeya, K., Kakioka, R., Cech, J.N., Osada, N., Leal, M.C., Inoue, J., Kume, M., Toyoda, A., et al. (2019). A key metabolic gene for recurrent freshwater colonization and radiation in fishes. Science 364, 886–889. 10.1126/science.aau5656.

13. Patterson, E.L., Pettinga, D.J., Ravet, K., Neve, P., and Gaines, T.A. (2018). Glyphosate resistance and *EPSPS* gene duplication: convergent evolution in multiple plant species. J. Hered. 109, 117–125. 10.1093/jhered/esx087.

14. Nguyen, D.-Q., Webber, C., Hehir-Kwa, J., Pfundt, R., Veltman, J., and Ponting, C.P. (2008). Reduced purifying selection prevails over positive selection in human copy number variant evolution. Genome Res. 18, 1711–1723. 10.1101/gr.077289.108.

15. Roudnitzky, N., Risso, D., Drayna, D., Behrens, M., Meyerhof, W., and Wooding, S.P. (2016). Copy number variation in *TAS2R* bitter taste receptor genes: structure, origin, and population genetics. Chem. Senses 41, 649–659. 10.1093/chemse/bjw067.

16. Endler, J. (1986). Natural selection in the wild (Princeton University Press).

17. Schluter, D. (2000). The ecology of adaptive radiation (Oxford University Press).

18. Bell, M.A., and Foster, S.A. (1994). The evolutionary biology of the threespine stickleback (Oxford University Press).

19. Taylor, E.B., and McPhail, J.D. (1986). Prolonged and burst swimming in anadromous and freshwater threespine stickleback, *Gasterosteus aculeatus*. Can. J. Zool. 64, 416–420. 10.1139/z86-064.

20. Reyes, M., and Baker, J.A. (2016). Prolonged swimming performance within the threespine stickleback (*Gasterosteus aculeatus*) adaptive radiation and the effect of dietary restriction. Evol. Ecol. Res. 17, 535–549.

21. Colosimo, P.F., Hosemann, K.E., Balabhadra, S., Jr, G.V., Dickson, M., Grimwood, J., Schmutz, J., Myers, R.M., Schluter, D., and Kingsley, D.M. (2005). Widespread parallel evolution in sticklebacks by repeated fixation of Ectodysplasin alleles. Science 307, 1928–1933. 10.1126/science.1107239.

22. Miller, C.T., Beleza, S., Pollen, A.A., Schluter, D., Kittles, R.A., Shriver, M.D., and Kingsley, D.M. (2007). *cis*-Regulatory changes in *Kit ligand* expression and parallel evolution of pigmentation in sticklebacks and humans. Cell 131, 1179–1189. 10.1016/j.cell.2007.10.055.

23. Jones, F.C., Chan, Y.F., Schmutz, J., Grimwood, J., Brady, S.D., Southwick, A.M., Absher, D.M., Myers, R.M., Reimchen, T.E., Deagle, B.E., et al. (2012). A genome-wide SNP genotyping array reveals patterns of global and repeated species-pair divergence in sticklebacks. Curr. Biol. 22, 83–90. 10.1016/j.cub.2011.11.045.

24. Hohenlohe, P.A., Bassham, S., Etter, P.D., Stiffler, N., Johnson, E.A., and Cresko, W.A. (2010). Population genomics of parallel adaptation in threespine stickleback using sequenced RAD tags. PLOS Genet. 6, e1000862. 10.1371/journal.pgen.1000862.

25. Jones, F.C., Grabherr, M.G., Chan, Y.F., Russell, P., Mauceli, E., Johnson, J., Swofford, R., Pirun, M., Zody, M.C., White, S., et al. (2012). The genomic basis of adaptive evolution in threespine sticklebacks. Nature 484, 55–61. 10.1038/nature10944.

26. Roberts Kingman, G.A., Vyas, D.N., Jones, F.C., Brady, S.D., Chen, H.I., Reid, K., Milhaven, M., Bertino, T.S., Aguirre, W.E., Heins, D.C., et al. (2021). Predicting future from past: The genomic basis of recurrent and rapid stickleback evolution. Sci. Adv. 7, eabg5285. 10.1126/sciadv.abg5285.

27. Chain, F.J.J., Feulner, P.G.D., Panchal, M., Eizaguirre, C., Samonte, I.E., Kalbe, M., Lenz, T.L., Stoll, M., Bornberg-Bauer, E., Milinski, M., et al. (2014). Extensive copy-number variation of young genes across stickleback populations. PLoS Genet. 10, e1004830. 10.1371/journal.pgen.1004830.

28. Hirase, S., Ozaki, H., and Iwasaki, W. (2014). Parallel selection on gene copy number variations through evolution of three-spined stickleback genomes. BMC Genomics 15, 735. 10.1186/1471-2164-15-735.

29. Lowe, C.B., Sanchez-Luege, N., Howes, T.R., Brady, S.D., Daugherty, R.R., Jones, F.C., Bell, M.A., and Kingsley, D.M. (2018). Detecting differential copy number variation between groups of samples. Genome Res. 28, 256–265. 10.1101/gr.206938.116.

30. Ikeda, D., Ono, Y., Snell, P., Edwards, Y.J.K., Elgar, G., and Watabe, S. (2007). Divergent evolution of the myosin heavy chain gene family in fish and tetrapods: Evidence from comparative genomic analysis. Physiol. Genomics 32, 1–15. 10.1152/physiolgenomics.00278.2006.

31. Ikeda, D., Ono, Y., Hirano, S., Kan-no, N., and Watabe, S. (2013). Lampreys have a single gene cluster for the fast skeletal myosin heavy chain gene family. PLOS ONE 8, e85500. 10.1371/journal.pone.0085500.

32. McGuigan, K., Phillips, P.C., and Postlethwait, J.H. (2004). Evolution of sarcomeric myosin heavy chain genes: Evidence from fish. Mol. Biol. Evol. 21, 1042–1056. 10.1093/molbev/msh103.

33. Chan, Y.F. (2009). The genomic basis of parallel evolution in three-spined stickleback (*Gasterosteus aculeatus*) (Doctoral dissertation, Stanford University).

34. Lee, L.A., Karabina, A., Broadwell, L.J., and Leinwand, L.A. (2019). The ancient sarcomeric myosins found in specialized muscles. Skelet. Muscle 9, 7. 10.1186/s13395-019-0192-3.

35. Ono, Y., Kinoshita, S., Ikeda, D., and Watabe, S. (2010). Early development of medaka *Oryzias latipes* muscles as revealed by transgenic approaches using embryonic and larval types of myosin heavy chain genes. Dev. Dyn. 239, 1807–1817. 10.1002/dvdy.22298.

36. Ono, Y., Liang, C., Ikeda, D., and Watabe, S. (2006). cDNA cloning of myosin heavy chain genes from medaka *Oryzias latipes* embryos and larvae and their expression patterns during development. Dev. Dyn. 235, 3092–3101. 10.1002/dvdy.20942.

37. Altschul, S.F., Gish, W., Miller, W., Myers, E.W., and Lipman, D.J. (1990). Basic local alignment search tool. J. Mol. Biol. 215, 403–410. 10.1016/S0022-2836(05)80360-2.

38. Nath, S., Shaw, D.E., and White, M.A. (2021). Improved contiguity of the threespine stickleback genome using long-read sequencing. G3 11, jkab007. 10.1093/g3journal/jkab007.

39. Hagen, D.W. (1967). Isolating mechanisms in threespine sticklebacks (*Gasterosteus*). J. Fish. Res. Bd. Can. 24, 1637–1692. 10.1139/f67-138.

40. McPhail, J.D. (1994). Speciation and the evolution of reproductive isolation in the sticklebacks (Gasterosteus) of south-western British Columbia. In The Evolutionary Biology of the Threespine Stickleback, M. A. Bell and S. a. Foster, eds. (Oxford University Press), pp. 399–437. 10.1093/oso/9780198577287.003.0014.

41. Jones, F.C., Brown, C., Pemberton, J.M., and Braithwaite, V.A. (2006). Reproductive isolation in a threespine stickleback hybrid zone. J. Evol. Biol. 19, 1531–1544. 10.1111/j.1420-9101.2006.01122.x.

42. Vines, T.H., Dalziel, A.C., Albert, A.Y.K., Veen, T., Schulte, P.M., and Schluter, D. (2016). Cline coupling and uncoupling in a stickleback hybrid zone. Evolution 70, 1023–1038. 10.1111/evo.12917.

43. Bell, M.A., Heins, D.C., Wund, M.A., Von Hippel, F.A., Massengill, R., Dunker, K., Bristow, G.A., and Aguirre, W.E. (2016). Reintroduction of threespine stickleback into Cheney and Scout Lakes, Alaska. Evol. Ecol. Res. 17, 157–178.

44. Bell, M.A., Aguirre, W.E., and Buck, N.J. (2004). Twelve years of contemporary armor evolution in a threespine stickleback population. Evolution 58, 814–824. 10.1111/j.0014-3820.2004.tb00414.x.

45. Kingsley, D.M., Zhu, B., Osoegawa, K., De Jong, P.J., Schein, J., Marra, M., Peichel, C., Amemiya, C., Schluter, D., Balabhadra, S., et al. (2004). New genomic tools for molecular studies of evolutionary change in threespine sticklebacks. Behaviour 141, 1331–1344.

46. Au, E.H., Weaver, S., Katikaneni, A., Wucherpfennig, J.I., Luo, Y., Mangan, R.J., Wund, M.A., Bell, M.A., and Lowe, C.B. (2025). Genome sequence of a marine threespine stickleback (*Gasterosteus aculeatus*) from Rabbit Slough in the Cook Inlet. G3 15, jkaf114. 10.1093/g3journal/jkaf114.

47. Wucherpfennig, J.I., Howes, T.R., Au, J.N., Au, E.H., Roberts Kingman, G.A., Brady, S.D., Herbert, A.L., Reimchen, T.E., Bell, M.A., Lowe, C.B., et al. (2022). Evolution of stickleback spines through independent *cis*-regulatory changes at *HOXDB*. Nat. Ecol. Evol. 6, 1537–1552. 10.1038/s41559-022-01855-3.

48. Ohno, S. (1970). Evolution by gene duplication (Springer).

49. Peichel, C.L., Ross, J.A., Matson, C.K., Dickson, M., Grimwood, J., Schmutz, J., Myers, R.M., Mori, S., Schluter, D., and Kingsley, D.M. (2004). The master sex-determination locus in threespine sticklebacks is on a nascent Y chromosome. Curr. Biol. 14, 1416–1424. 10.1016/j.cub.2004.08.030.

50. Peichel, C.L., McCann, S.R., Ross, J.A., Naftaly, A.F.S., Urton, J.R., Cech, J.N., Grimwood, J., Schmutz, J., Myers, R.M., Kingsley, D.M., et al. (2020). Assembly of the threespine stickleback Y chromosome reveals convergent signatures of sex chromosome evolution. Genome Biol. 21, 177. 10.1186/s13059-020-02097-x.

51. Birchler, J.A., and Yang, H. (2022). The multiple fates of gene duplications: Deletion, hypofunctionalization, subfunctionalization, neofunctionalization, dosage balance constraints, and neutral variation. Plant Cell 34, 2466–2474. 10.1093/plcell/koac076.

52. Loehlin, D.W., and Carroll, S.B. (2016). Expression of tandem gene duplicates is often greater than twofold. Proc. Natl. Acad. Sci. U.S.A. 113, 5988–5992. 10.1073/pnas.1605886113.

53. Huang, K.M., and Chain, F.J.J. (2021). Copy number variations and young duplicate genes have high methylation levels in sticklebacks. Evolution 75, 706–718. 10.1111/evo.14184.

54. Rogers, R.L., Shao, L., and Thornton, K.R. (2017). Tandem duplications lead to novel expression patterns through exon shuffling in *Drosophila yakuba*. PLOS Genet. 13, e1006795. 10.1371/journal.pgen.1006795.

55. Hastings, P.J., Lupski, J.R., Rosenberg, S.M., and Ira, G. (2009). Mechanisms of change in gene copy number. Nat. Rev. Genet. 10, 551–564. 10.1038/nrg2593.

56. Stankiewicz, P., and Lupski, J.R. (2002). Genome architecture, rearrangements and genomic disorders. TIG 18, 74–82. 10.1016/S0168-9525(02)02592-1.

57. Hastings, P.J., Ira, G., and Lupski, J.R. (2009). A microhomology-mediated break-induced replication model for the origin of human copy number variation. PLOS Genet. 5, e1000327. 10.1371/journal.pgen.1000327.

58. Lowey, S., Saraswat, L.D., Liu, H., Volkmann, N., and Hanein, D. (2007). Evidence for an interaction between the SH3 domain and the N-terminal extension of the essential light chain in class II myosins. J. Mol. Biol. 371, 902–913. 10.1016/j.jmb.2007.05.080.

59. Abramson, J., Adler, J., Dunger, J., Evans, R., Green, T., Pritzel, A., Ronneberger, O., Willmore, L., Ballard, A.J., Bambrick, J., et al. (2024). Accurate structure prediction of biomolecular interactions with AlphaFold 3. Nature 630, 493–500. 10.1038/s41586-024-07487-w.

60. Papp, B., Pál, C., and Hurst, L.D. (2003). Dosage sensitivity and the evolution of gene families in yeast. Nature 424, 194–197. 10.1038/nature01771.

61. Konrad, A., Flibotte, S., Taylor, J., Waterston, R.H., Moerman, D.G., Bergthorsson, U., and Katju, V. (2018). Mutational and transcriptional landscape of spontaneous gene duplications and deletions in *Caenorhabditis elegans*. Proc. Natl. Acad. Sci. U.S.A. 115, 7386–7391. 10.1073/pnas.1801930115.

62. Watabe, S. (2002). Temperature plasticity of contractile proteins in fish muscle. J. Exp. Biol. 205, 2231–2236. 10.1242/jeb.205.15.2231.

63. Uyeda, T.Q.P., Ruppel, K.M., and Spudich, J.A. (1994). Enzymatic activities correlate with chimaeric substitutions at the actin-binding face of myosin. Nature 368, 567–569. 10.1038/368567a0.

64. Moss, R.L., Diffee, G.M., and Greaser, M.L. (1995). Contractile properties of skeletal muscle fibers in relation to myofibrillar protein isoforms. In Reviews of Physiology, Biochemistry and Pharmacology, (Springer). 10.1007/BFb0049775.

65. Goodson, H.V., Warrick, H.M., and Spudich, J.A. (1999). Specialized conservation of surface loops of myosin: evidence that loops are involved in determining functional characteristics. J. Mol. Biol. 287, 173–185. 10.1006/jmbi.1999.2565.

66. Dalziel, A.C., Ou, M., and Schulte, P.M. (2012). Mechanisms underlying parallel reductions in aerobic capacity in non-migratory threespine stickleback (*Gasterosteus aculeatus*) populations. J. Exp. Biol. 215, 746–759. 10.1242/jeb.065425.

67. te Kronnie, G., Tatarczuch, L., van Raamsdonk, W., and Kilarski, W. (1983). Muscle fibre types in the myotome of stickleback, *Gasterosteus aculeatus* L.; a histochemical, immunohistochemical and ultrastructural study. J. Fish Biol. 22, 303–316. 10.1111/j.1095-8649.1983.tb04754.x.

68. Garcia de la serrana, D., Wreggelsworth, K., and Johnston, I.A. (2018). Duplication of a single *myhz1.1* gene facilitated the ability of goldfish (*Carassius auratus*) to alter fast muscle contractile properties with seasonal temperature change. Front. Physiol. 9, 1724. 10.3389/fphys.2018.01724.

69. Südhof, T.C. (2002). Synaptotagmins: Why so many? J. Biol. Chem. 277, 7629–7632. 10.1074/jbc.R100052200.

70. Gustavsson, N., and Han, W. (2009). Calcium-sensing beyond neurotransmitters: functions of synaptotagmins in neuroendocrine and endocrine secretion. Biosci. Rep. 29, 245–259. 10.1042/BSR20090031.

71. Bhoyar, R.C., Jadhao, A.G., Sabharwal, A., Ranjan, G., Sivasubbu, S., and Pinelli, C. (2019). Knockdown of calcium-binding *calb2a* and *calb2b* genes indicates the key regulator of the early development of the zebrafish, *Danio rerio*. Brain Struct. Funct. 224, 627–642. 10.1007/s00429-018-1797-8.

72. Wu, X.M., Hu, Y.W., Xue, N.N., Ren, S.S., Chen, S.N., Nie, P., and Chang, M.X. (2017). Role of zebrafish NLRC5 in antiviral response and transcriptional regulation of MHC related genes. Dev. Comp. Immunol. 68, 58–68. 10.1016/j.dci.2016.11.018.

73. Cao, L., Wu, X.M., Hu, Y.W., Xue, N.N., Nie, P., and Chang, M.X. (2018). The discrepancy function of NLRC5 isoforms in antiviral and antibacterial immune responses. Dev. Comp. Immunol. 84, 153–163. 10.1016/j.dci.2018.02.013.

74. Benkő, S., Kovács, E.G., Hezel, F., and Kufer, T.A. (2017). NLRC5 Functions beyond MHC I Regulation—What Do We Know So Far? Front. Immunol. 8. 10.3389/fimmu.2017.00150.

75. Kapitonov, V.V., and Jurka, J. (2004). *Harbinger* transposons and an ancient HARBI1 gene derived from a transposase. DNA Cell Biol. 23, 311–324. 10.1089/104454904323090949.

76. Sun, B., Qian, X., and Zhu, F. (2018). Molecular characterization of shrimp harbinger transposase derived 1 (HARBI1)-like and its role in white spot syndrome virus and *Vibrio alginolyticus* infection. Fish Shellfish Immunol. 78, 222–232. 10.1016/j.fsi.2018.04.032.

77. Reimchen, T.E., Steeves, D., and Bergstrom, C.A. (2016). Sex matters for defence and trophic traits of threespine stickleback. Evol. Ecol. Res. 17, 459–485.

78. Kitano, J., Kakioka, R., Ishikawa, A., Toyoda, A., and Kusakabe, M. (2020). Differences in the contributions of sex linkage and androgen regulation to sex-biased gene expression in juvenile and adult sticklebacks. J. Evol. Biol. 33, 1129–1138. 10.1111/jeb.13662.

79. Schutz, H., Anderson, R.J., Warwick, E.G., Barry, T.N., and Jamniczky, H.A. (2022). Sexually mediated phenotypic variation within and between sexes as a continuum structured by ecology: The mosaic nature of skeletal variation across body regions in Threespine stickleback (*Gasterosteus aculeatus* L.). Ecol. Evol. 12, e9367. 10.1002/ece3.9367.

80. Blain, S.A., Roesti, M., Thompson, K.A., Kinney, M.H., and Schluter, D. (2025). Species interactions, divergence, and the rapid evolution of ecological sexual dimorphism in threespine sticklebacks. Preprint at bioRxiv. 10.1101/2025.10.26.684666.

81. Martin, A., and Orgogozo, V. (2013). The loci of repeated evolution: A catalog of genetic hotspots of phenotypic variation. Evolution 67, 1235–1250. 10.1111/evo.12081.

82. Schluter, D., and Conte, G.L. (2009). Genetics and ecological speciation. Proc. Natl. Acad. Sci. U.S.A. 106, 9955–9962. 10.1073/pnas.0901264106.

83. Kitano, J., Lema, S.C., Luckenbach, J.A., Mori, S., Kawagishi, Y., Kusakabe, M., Swanson, P., and Peichel, C.L. (2010). Adaptive divergence in the thyroid hormone signaling pathway in the stickleback radiation. Curr. Biol. 20, 2124–2130. 10.1016/j.cub.2010.10.050.

84. Chan, Y.F., Marks, M.E., Jones, F.C., Villarreal, G., Shapiro, M.D., Brady, S.D., Southwick, A.M., Absher, D.M., Grimwood, J., Schmutz, J., et al. (2010). Adaptive evolution of pelvic reduction in sticklebacks by recurrent deletion of a *Pitx1* enhancer. Science 327, 302–305. 10.1126/science.1182213.

85. Xie, K.T., Wang, G., Thompson, A.C., Wucherpfennig, J.I., Reimchen, T.E., MacColl, A.D.C., Schluter, D., Bell, M.A., Vasquez, K.M., and Kingsley, D.M. (2019). DNA fragility in the parallel evolution of pelvic reduction in stickleback fish. Science 363, 81–84. 10.1126/science.aan1425.

86. Deagle, B.E., Jones, F.C., Chan, Y.F., Absher, D.M., Kingsley, D.M., and Reimchen, T.E. (2012). Population genomics of parallel phenotypic evolution in stickleback across stream-lake ecological transitions. Proc. Biol. Sci. 279, 1277–1286. 10.1098/rspb.2011.1552.

87. Kawakami, K. (2007). *Tol2*: a versatile gene transfer vector in vertebrates. Genome Biol. 8, S7. 10.1186/gb-2007-8-s1-s7.

88. Katoh, K., and Standley, D.M. (2013). MAFFT multiple sequence alignment software version 7: improvements in performance and usability. Mol. Biol. Evol. 30, 772–780. 10.1093/molbev/mst010.

89. Katoh, K., Misawa, K., Kuma, K., and Miyata, T. (2002). MAFFT: a novel method for rapid multiple sequence alignment based on fast Fourier transform. Nucleic Acids Res. 30, 3059–3066. 10.1093/nar/gkf436.

90. Quinlan, A.R., and Hall, I.M. (2010). BEDTools: a flexible suite of utilities for comparing genomic features. Bioinformatics 26, 841–842. 10.1093/bioinformatics/btq033.

91. Lopez-Delisle, L., Rabbani, L., Wolff, J., Bhardwaj, V., Backofen, R., Grüning, B., Ramírez, F., and Manke, T. (2021). pyGenomeTracks: reproducible plots for multivariate genomic datasets. Bioinformatics 37, 422–423. 10.1093/bioinformatics/btaa692.

92. Danecek, P., Bonfield, J.K., Liddle, J., Marshall, J., Ohan, V., Pollard, M.O., Whitwham, A., Keane, T., McCarthy, S.A., Davies, R.M., et al. (2021). Twelve years of SAMtools and BCFtools. GigaScience 10, giab008. 10.1093/gigascience/giab008.

93. Li, H., and Durbin, R. (2009). Fast and accurate short read alignment with Burrows–Wheeler transform. Bioinformatics 25, 1754–1760. 10.1093/bioinformatics/btp324.

94. Huang, W., Li, L., Myers, J.R., and Marth, G.T. (2012). ART: a next-generation sequencing read simulator. Bioinformatics 28, 593–594. 10.1093/bioinformatics/btr708.

95. Minh, B.Q., Schmidt, H.A., Chernomor, O., Schrempf, D., Woodhams, M.D., von Haeseler, A., and Lanfear, R. (2020). IQ-TREE 2: New models and efficient methods for phylogenetic inference in the genomic era. Mol. Biol. Evol. 37, 1530–1534. 10.1093/molbev/msaa015.

96. Wong, T.K.F., Ly-Trong, N., Ren, H., Baños, H., Roger, A.J., Susko, E., Bielow, C., De Maio, N., Goldman, N., Hahn, M.W., et al. (2025). IQ-TREE 3: Phylogenomic inference software using complex evolutionary models. Preprint at EcoEvoRxiv. 10.32942/X2P62N.

97. Kalyaanamoorthy, S., Minh, B.Q., Wong, T.K.F., von Haeseler, A., and Jermiin, L.S. (2017). ModelFinder: fast model selection for accurate phylogenetic estimates. Nat. Methods 14, 587–589. 10.1038/nmeth.4285.

98. Yu, G., Smith, D.K., Zhu, H., Guan, Y., and Lam, T.T.-Y. (2017). ggtree: an R package for visualization and annotation of phylogenetic trees with their covariates and other associated data. Methods Ecol. Evol. 8, 28–36. 10.1111/2041-210X.12628.

99. Cheng, H., Concepcion, G.T., Feng, X., Zhang, H., and Li, H. (2021). Haplotype-resolved *de novo* assembly using phased assembly graphs with hifiasm. Nat. Methods 18, 170–175. 10.1038/s41592-020-01056-5.

100. Koren, S., Walenz, B.P., Berlin, K., Miller, J.R., Bergman, N.H., and Phillippy, A.M. (2017). Canu: scalable and accurate long-read assembly via adaptive *k*-mer weighting and repeat separation. Genome Res. 27, 722–736. 10.1101/gr.215087.116.

101. Kent, W.J. (2002). BLAT—The BLAST-like alignment tool. Genome Res. 12, 656–664. 10.1101/gr.229202.

102. Shumate, A., and Salzberg, S.L. (2021). Liftoff: accurate mapping of gene annotations. Bioinformatics 37, 1639–1643. 10.1093/bioinformatics/btaa1016.

103. Li, H. (2018). Minimap2: pairwise alignment for nucleotide sequences. Bioinformatics 34, 3094–3100. 10.1093/bioinformatics/bty191.

104. Bianchini, G., and Sánchez-Baracaldo, P. (2024). TreeViewer: Flexible, modular software to visualise and manipulate phylogenetic trees. Ecol. Evol. 14, e10873. 10.1002/ece3.10873.

105. Blum, M., Andreeva, A., Florentino, L.C., Chuguransky, S.R., Grego, T., Hobbs, E., Pinto, B.L., Orr, A., Paysan-Lafosse, T., Ponamareva, I., et al. (2025). InterPro: the protein sequence classification resource in 2025. Nucleic Acids Res. 53, D444–D456. 10.1093/nar/gkae1082.

106. Chen, S., Zhou, Y., Chen, Y., and Gu, J. (2018). fastp: an ultra-fast all-in-one FASTQ preprocessor. Bioinformatics 34, i884–i890. 10.1093/bioinformatics/bty560.

107. Broman, K.W., Wu, H., Sen, Ś., and Churchill, G.A. (2003). R/qtl: QTL mapping in experimental crosses. Bioinformatics 19, 889–890. 10.1093/bioinformatics/btg112.

108. Hinrichs, A.S., Karolchik, D., Baertsch, R., Barber, G.P., Bejerano, G., Clawson, H., Diekhans, M., Furey, T.S., Harte, R.A., Hsu, F., et al. (2006). The UCSC Genome Browser Database: update 2006. Nucleic Acids Res. 34, D590–D598. 10.1093/nar/gkj144.

109. Broad Institute (2023). Picard Toolkit (version 3.1.1). https://broadinstitute.github.io/picard/.

110. Kassambara, A. (2025). ggpubr: “ggplot2” based publication ready plots. https://rpkgs.datanovia.com/ggpubr/.

111. Oritz, E.M. (2019). vcf2phylip v2.0: convert a VCF matrix into several matrix formats for phylogenetic analysis. 10.5281/zenodo.2540861.

112. Pohlert, T. (2023). trend: Non-parametric trend tests and change-point detection. https://CRAN.R-project.org/package=trend.

113. Velden, N. van der (2025). geneviewer: Gene cluster visualizations. https://github.com/nvelden/geneviewer.

114. Livak, K.J., and Schmittgen, T.D. (2001). Analysis of relative gene expression data using real-time quantitative PCR and the 2^−ΔΔCT^ method. Methods 25, 402–408. 10.1006/meth.2001.1262.

115. Wickham, H. (2016). ggplot2: Elegant Graphics for Data Analysis. https://ggplot2.tidyverse.org.

116. Babraham Bioinformatics (2023). FastQC: a quality control tool for high throughput sequence Data. https://www.bioinformatics.babraham.ac.uk/projects/fastqc/.

117. ImageMagick Studio LLC (2024). ImageMagick. https://imagemagick.org/.

