## Supplementary material for "Rapid and repeated evolution of myosin copy number in threespine stickleback": DocumentS1

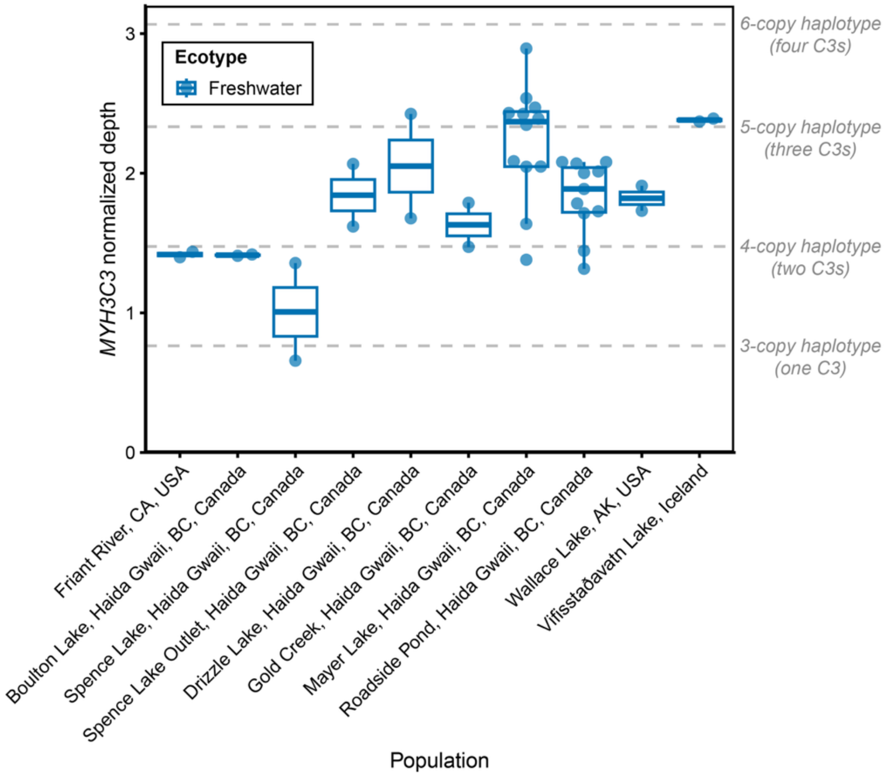


**Figure S1. Within-population variation in *MYH3C3* copy number, related to Figure 1**

*MYH3C3* read depth normalized by mean autosomal read depth determined for female stickleback from Roberts Kingman et al.[S1] in populations represented by at least two individual fish. Roadside Pond was derived from a transplant of Mayer Lake fish in 1993[2]. Read depth differences support variation in *MYH3C3* copy number within Drizzle Lake, Spence Lake, Mayer Lake, and Roadside Pond. Dashed lines indicate read depth calculated from simulated reads of different assembled 3- to 6-copy myosin haplotypes (one to four *MYH3C3* copies per haplotype; Figure 2). See Table S1 and Roberts Kingman et al.[S1] for more information about each sample.

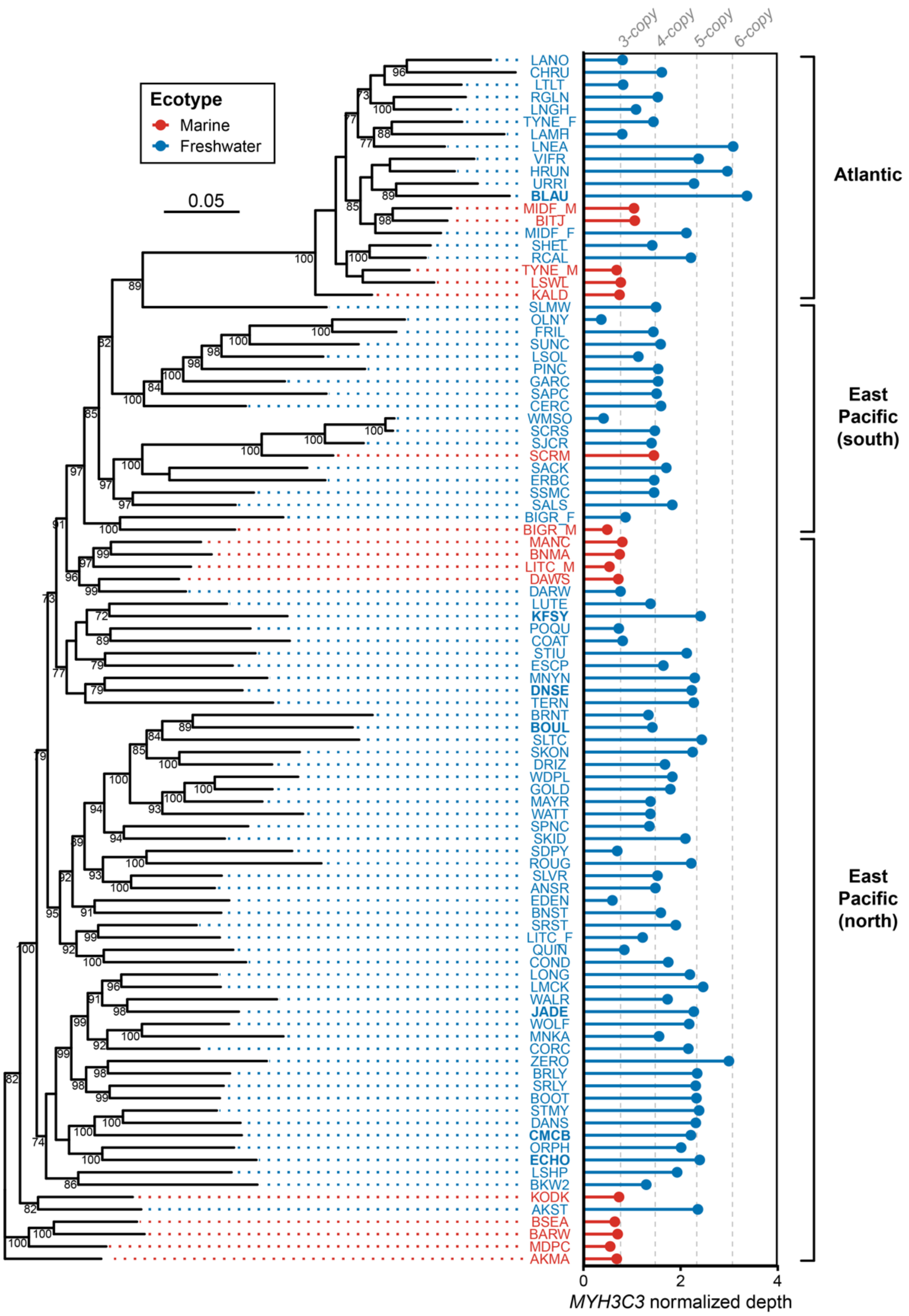


**Figure S2. Repeated evolution of *MYH3C3* copy number across global stickleback populations, related to Figure 1**

Phylogenetic tree based on 100,000 genome-wide neutral SNPs from the 15 marine (red) and 81 freshwater (blue) stickleback from Figure 1B. Bootstrap values of ≥70 are labeled below each node. Branch lengths are based on the number of inferred substitutions, indicated by the scale bar. Normalized *MYH3C3* read depth is plotted for each individual as previously described in Figure 1B. Freshwater populations with higher *MYH3C3* read depth are interspersed with marine populations with lower *MYH3C3* read depth in both the Pacific and Atlantic basins, suggesting *MYH3C3* copy number expansions have likely evolved multiple times. See Table S1 and Roberts Kingman et al.[S1] for more information about each sample.

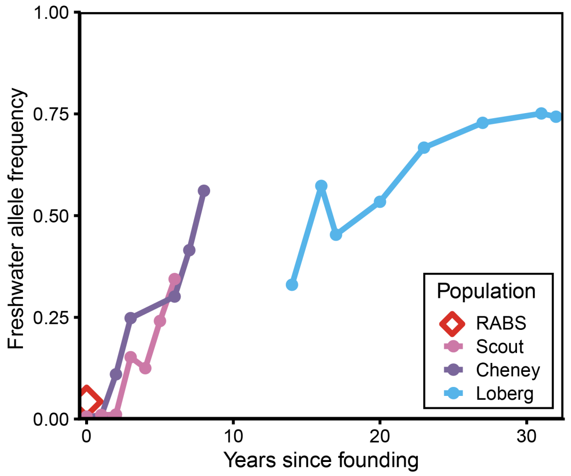


**Figure S3. SNP allele frequencies associated with the freshwater *MYH3C* allele increase rapidly when marine fish are introduced to freshwater, related to Figure 1**

Allele frequencies for the most significant SNP (ChrXIX:2,744,769 [*gasAcu1-4* reference]) within the Sensitive TempoPeak[S1] overlapping the ChrXIX *MYH3C* locus. Allele frequencies of the freshwater allele are low in the founder marine population (RABS, red) but rise to over 30% frequency within several generations in freshwater habitats (Scout [pink], Cheney [purple], Loberg [light blue]).

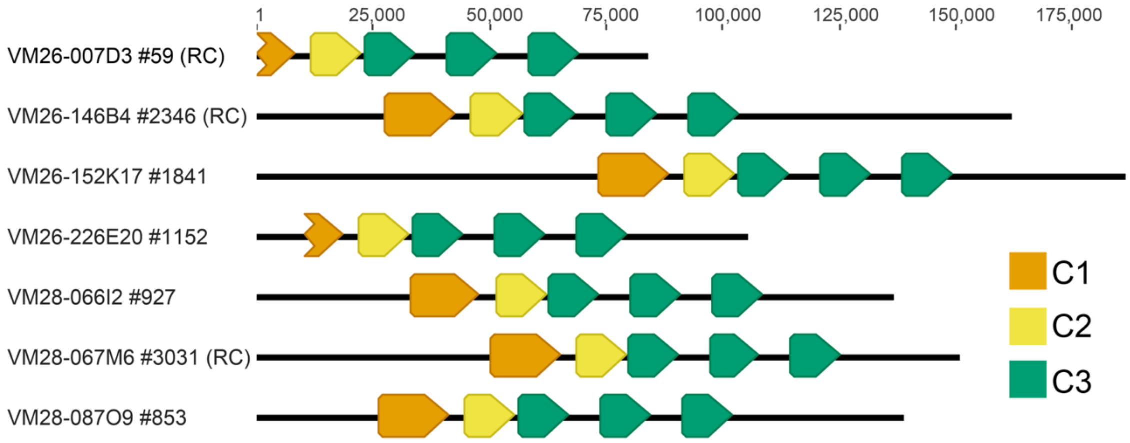


**Figure S4. The BEPA reference stickleback has a 5-copy *MYH3C* haplotype, related to Figure 1 and Figure 2**

Representative individual Nanopore reads from all seven BACs that fully span duplicated *C3* copies derived from the female BEPA stickleback used to create the freshwater reference genome[S3,S4]. Individual DNA molecules were annotated with *C1* (orange), *C2* (yellow), and *C3* (green). All seven clones show three *C3* gene copies at the myosin locus, suggesting that the freshwater reference genome artificially collapsed *C3* copies during assembly. Some reads have been reverse complemented for visual clarity (RC). The reads from the VM26-007D3 and VM26-226E20 BAC clones only partially span *C1*, due the lengths of the longest *C3* spanning read (VM26-007D3) or to the size of the stickleback insert (VM26-226E20).

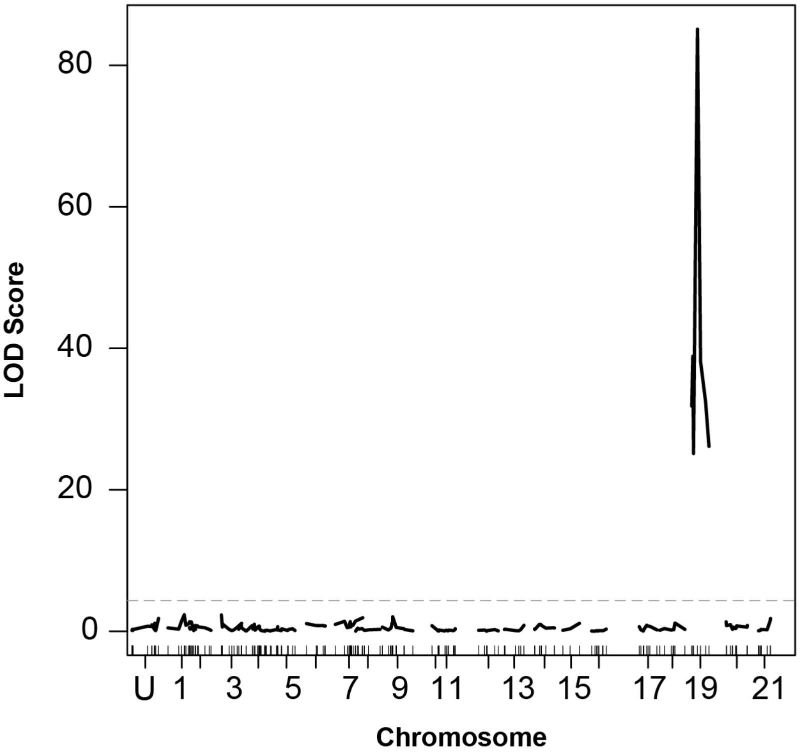


**Figure S5. Myosin *C3* copy numbers map to a single major locus, related to Figure 2**

*C3* copy numbers were scored in 440 individual F2 progeny from a cross between marine (Bodega Bay, CA) and freshwater (Boulton Lake, BC, Canada) stickleback. A SNP marker adjacent to the *MYH3C* locus on ChrXIX explained nearly 60% of the variance in copy number (LOD = 85.1, PVE = 0.59, $p < 1x10^{-4}$), and no significant linkage was found to other chromosomes. The LOD score cutoff for a genome-wide $\alpha = 0.01$ is shown with the dashed line. ChrU represents concatenated unassembled regions of the genome[S3].

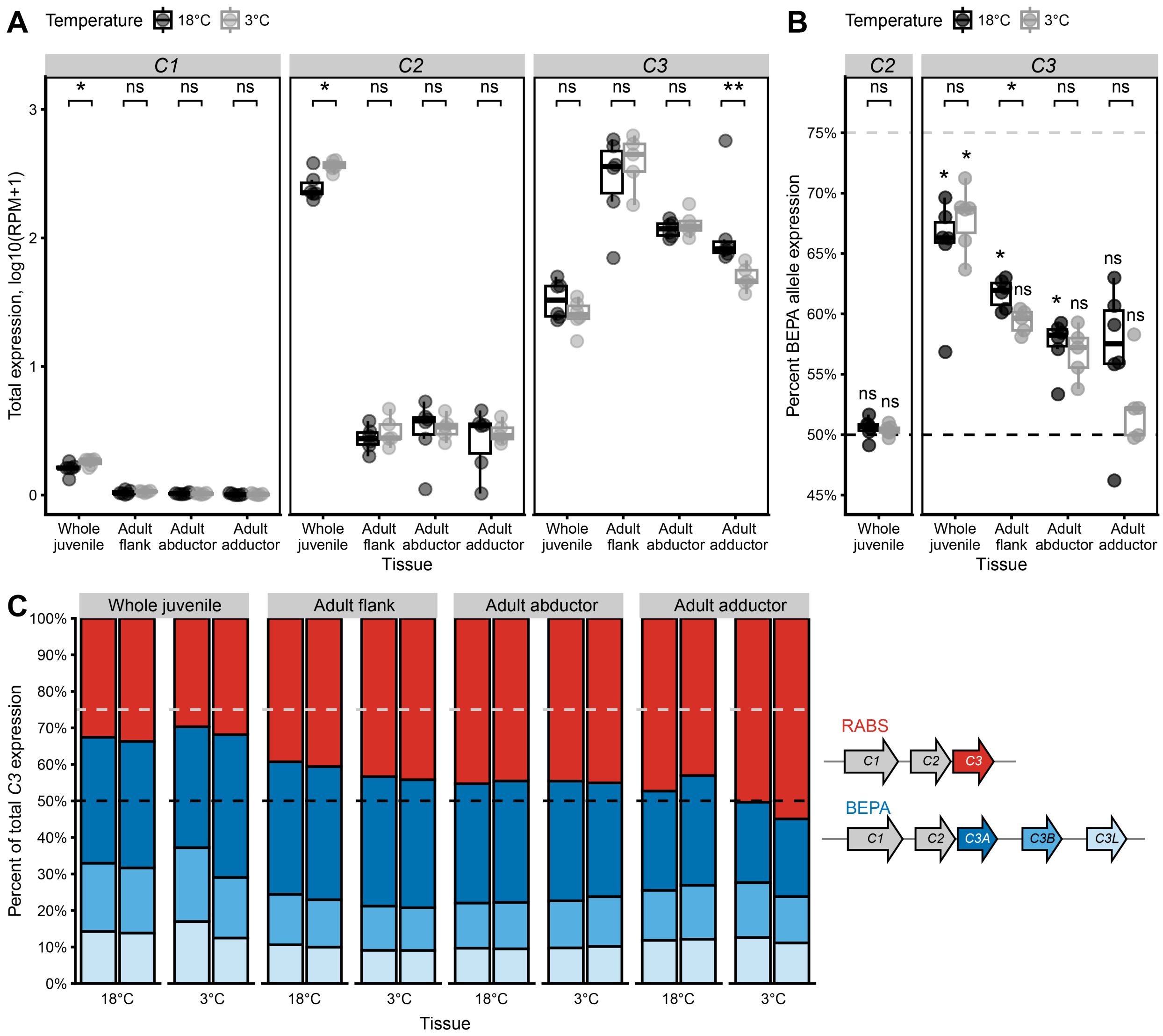


**Figure S6.** ***MYH3C*** **expression at different temperatures, related to Figure 3**

(A) Box plots showing the relative expression levels of *MYH3C* copies in RABS x BEPA F1 hybrid females based on seven sets of MCE 27-mers. Fish were acclimated to either standard 18°C (black) or cold 3°C (grey) temperatures. For all conditions, $n$ = 6, except for 3°C adult tissues where $n$ = 5. Gene expression differences under each temperature condition were compared using a two-sided Wilcoxon rank sum test. Expression of *C1* and *C2* increases at 3°C in juveniles while expression of *C3* decreases at 3°C in adult adductor muscle.

(B) Box plots showing allele-specific expression (ASE) of the BEPA vs. RABS alleles based on ASE *k*-mer sets (four *C2* sets and nine *C3* sets) for each *MYH3C* copy under 18°C or 3°C conditions. Deviation from an equal ratio of expression from the BEPA and RABS alleles (50% BEPA allele expression, black dashed line) was evaluated using a one-sample Wilcoxon rank sum test (mu = 0.5; shown directly above each box plot). The BEPA allele expression would be 75% with expression proportional to *C3* copy number (grey dashed line). Gene expression differences under each temperature condition were compared using a two-sided Wilcoxon rank sum test (shown above each bracket). Adult tissues at 3°C are underpowered for detecting significant ASE due to reduced sample number. Expression of *C3* from the BEPA allele decreases at 3°C in adult flank muscle.

(C) Stacked bar plots representing the relative expression levels of *C3* copies from RABS (red) and BEPA (*C3A*, dark blue; *C3B*, medium blue; *C3L*, light blue) determined by mapped Kinnex reads.

ns = not significant, $p > 0.05$, *$p \leq 0.05$, **$p \leq 0.01$

| Position | MYH3C copy | Marine aa | Freshwater aa | Marine aa type | Freshwater aa type | Same type? | Domain |
|---|---|---|---|---|---|---|---|
| 57 | C1 | A | T | hydrophobic | polar | no | SH3 |
| 209 | C1 | T | A | polar | hydrophobic | no | Myosin motor |
| 388 | C1 | M | L | hydrophobic | hydrophobic | yes | Myosin motor |
| 468 | C1 | Y | F | hydrophobic | hydrophobic | yes | Myosin motor |
| 470 | C1 | S | T | polar | polar | yes | Myosin motor |
| 471 | C1 | M | L | hydrophobic | hydrophobic | yes | Myosin motor |
| 507 | C1 | D | E | negative charge | negative charge | no | Myosin motor |
| 571 | C1 | A | V | hydrophobic | hydrophobic | yes | Myosin motor |
| 573 | C1 | A | G | hydrophobic | special | no | Myosin motor |
| 728 | C1 | D | E | negative charge | negative charge | no | Myosin motor |
| 1019 | C1 | A | S | hydrophobic | polar | no | Myosin tail |
| 1058 | C1 | A | S | hydrophobic | polar | no | Myosin tail |
| 1062 | C1 | I | V | hydrophobic | hydrophobic | yes | Myosin tail |
| 1233 | C1 | G | A | special | hydrophobic | no | Myosin tail |
| 1272 | C1 | H | Q | positive charge | polar | no | Myosin tail |
| 1859 | C1 | R | K | positive charge | positive charge | yes | Myosin tail |
| 29 | C2 | S | T | polar | polar | yes | - |
| 43 | C2 | A | V | hydrophobic | hydrophobic | yes | SH3 |
| 55 | C3 | K | R | positive charge | positive charge | yes | SH3 |
| 56 | C3 | E | D | negative charge | negative charge | no | SH3 |
| 76 | C3 | D | E | negative charge | negative charge | no | SH3 |
| 209 | C3 | T | A | polar | hydrophobic | no | Myosin motor |
| 251 | C3 | S | G | polar | special | no | Myosin motor |
| 257 | C3 | A | S | hydrophobic | polar | no | Myosin motor |
| 325 | C3 | I | V | hydrophobic | hydrophobic | yes | Myosin motor |
| 573 | C3 | G | A | special | hydrophobic | no | Myosin motor |
| 589 | C3 | T | N | polar | polar | yes | Myosin motor |
| 838 | C3 | T | S | polar | polar | yes | - |
| 885 | C3 | L | M | hydrophobic | hydrophobic | yes | Myosin tail |
| 1019 | C3 | A | S | hydrophobic | polar | no | Myosin tail |
| 1186 | C3 | S | A | polar | hydrophobic | no | Myosin tail |
| 1244 | C3 | L | M | hydrophobic | hydrophobic | yes | Myosin tail |
| 1365 | C3 | A | S | hydrophobic | polar | no | Myosin tail |
| 1608 | C3 | L | M | hydrophobic | hydrophobic | yes | Myosin tail |
| 1824 | C3 | A | T | hydrophobic | polar | no | Myosin tail |
| 1929 | C3 | A | V | hydrophobic | hydrophobic | yes | - |

**Table S3. Marine/freshwater divergent amino acids in each *MYH3C* copy, related to Figure 2**

| Set | *MYH3C* target | Allele target | Exon | *k*-mer (forward) | *k*-mer (reverse complement) |
|---|---|---|---|---|---|
| MCE1 | *C1* | both | 3 | GCTATTTACCTTCGTAAGCCAGAGAGG | CCTCTCTGGCTTACGAAGGTAAATAGC |
| MCE1 | *C2* | both | 3 | GCCATTTACCTTCGTAAGCCAGAGAAG | CTTCTCTGGCTTACGAAGGTAAATGGC |
| MCE1 | *C3* | both | 3 | GCCATTTACCTTCGTAAGCCAGAGAGG | CCTCTCTGGCTTACGAAGGTAAATGGC |
| MCE3 | *C1* | both | 3 | TGTACTTGAAGGCCACAATCCTCAAGA | TCTTGAGGATTGTGGCCTTCAAGTACA |
| MCE3 | *C2* | both | 3 | TGTACTTGAAGGCCACAATCATCAAGA | TCTTGATGATTGTGGCCTTCAAGTACA |
| MCE3 | *C3* | both | 3 | TGTATTTGAAGGCCAAAGTCATCAAGA | TCTTGATGACTTTGGCCTTCAAATACA |
| MCE5 | *C1* | both | 28 | AGGACCAACTTAGCGAAGTGAAGACAA | TTGTCTTCACTTCGCTAAGTTGGTCCT |
| MCE5 | *C2* | both | 28 | AGGACCAACTTAGCGAACTGAAGACAA | TTGTCTTCAGTTCGCTAAGTTGGTCCT |
| MCE5 | *C3* | both | 28 | AGGACCAATTTAGCGAACTGAAGACAA | TTGTCTTCAGTTCGCTAAATTGGTCCT |
| MCE6 | *C1* | both | 33 | GTGGAAACGGAGAAGGCTGAGATCCAG | CTGGATCTCAGCCTTCTCCGTTTCCAC |
| MCE6 | *C2* | both | 33 | GTGGAAACAGAGAAGTCTGAGATCCAG | CTGGATCTCAGACTTCTCTGTTTCCAC |
| MCE6 | *C3* | both | 33 | GTGGAAACAGAGAAGACTGAGATCCAG | CTGGATCTCAGTCTTCTCTGTTTCCAC |
| MCE7 | *C1* | both | 37 | GAGCAAGACACGAGTGCTCACCTTGAG | CTCAAGGTGAGCACTCGTGTCTTGCTC |
| MCE7 | *C2* | both | 37 | GAGCAAGACACTAGTGCTCACCTTGAG | CTCAAGGTGAGCACTAGTGTCTTGCTC |
| MCE7 | *C3* | both | 37 | GAGCAAGACACGAGTTCTCACCTTGAG | CTCAAGGTGAGAACTCGTGTCTTGCTC |
| MCE8 | *C1* | both | 40 | AATGCTCATCTGTCCAAATGCAGGAAG | CTTCCTGCATTTGGACAGATGAGCATT |
| MCE8 | *C2* | both | 40 | AATGCTCATCTGTCCAAGTGCAGGAAG | CTTCCTGCACTTGGACAGATGAGCATT |
| MCE8 | *C3* | both | 40 | AATACTCATCTGTCCAAGTGCAGAAAG | CTTTCTGCACTTGGACAGATGAGTATT |
| MCE9 | *C1* | both | 40 | AACAAGCTGAGAGCAAAAACCCGTGAC | GTCACGGGTTTTTGCTCTCAGCTTGTT |
| MCE9 | *C2* | both | 40 | AACAAGCTGAGAGCAAAAAGCCGTGAC | GTCACGGCTTTTTGCTCTCAGCTTGTT |
| MCE9 | *C3* | both | 40 | AACAAGATGAGAGCAAAAAGTCGTGAC | GTCACGACTTTTTGCTCTCATCTTGTT |
| C1_ASE3 | *C1* | BEPA | 3 | CCTCAAGAAAGAGACTGGCAAAGTCAC | GTGACTTTGCCAGTCTCTTTCTTGAGG |
| C1_ASE3 | *C1* | RABS | 3 | CCTCAAGAAAGAGGCTGGCAAAGTCAC | GTGACTTTGCCAGCCTCTTTCTTGAGG |
| C1_ASE5 | *C1* | BEPA | 12 | TTCTCAAGTGAAGAGAAGTTGAGCATC | GATGCTCAACTTCTCTTCACTTGAGAA |
| C1_ASE5 | *C1* | RABS | 12 | TTCTCAAGTGAAGAGAAGCTGAGCATC | GATGCTCAGCTTCTCTTCACTTGAGAA |
| C1_ASE6 | *C1* | BEPA | 21 | CACTGGTCACCATGACTCAGGCTTTGT | ACAAAGCCTGAGTCATGGTGACCAGTG |
| C1_ASE6 | *C1* | RABS | 21 | CACTGGTCACAATGACTCAGGCTTTGT | ACAAAGCCTGAGTCATTGTGACCAGTG |
| C1_ASE7 | *C1* | BEPA | 27 | GATGGAAATTGATGACCTCTCTAGCAA | TTGCTAGAGAGGTCATCAATTTCCATC |
| C1_ASE7 | *C1* | RABS | 27 | GATGGAAATCGATGACCTCTCCAGCAA | TTGCTGGAGAGGTCATCGATTTCCATC |
| C1_ASE8 | *C1* | BEPA | 27 | CCTCTCTAGCAACATGGAGGCTGTTGC | GCAACAGCCTCCATGTTGCTAGAGAGG |
| C1_ASE8 | *C1* | RABS | 27 | CCTCTCCAGCAACATGGAGGGTGTTGC | GCAACACCCTCCATGTTGCTGGAGAGG |
| C1_ASE9 | *C1* | BEPA | 36 | AGCTTGAGACTGACCTGGTCCAAGTCC | GGACTTGGACCAGGTCAGTCTCAAGCT |
| C1_ASE9 | *C1* | RABS | 36 | AGCTCGAGACTGACCTGGTCCAGGTCC | GGACCTGGACCAGGTCAGTCTCGAGCT |
| C1_ASE10 | *C1* | BEPA | 38 | AGCAAAGACGTGGAGCAGATGCCGTTA | TAACGGCATCTGCTCCACGTCTTTGCT |
| C1_ASE10 | *C1* | RABS | 38 | AGCAGAGACGTGGAGCAGATGCCGTTA | TAACGGCATCTGCTCCACGTCTCTGCT |
| C1_ASE11 | *C1* | BEPA | 41 | AGTGTATGTGTTAAGTCATATAATATG | CATATTATATGACTTAACACATACACT |
| C1_ASE11 | *C1* | RABS | 41 | AGTGTATGTGTTAAATCATATAATATG | CATATTATATGATTTAACACATACACT |
| C2_ASE1 | *C2* | BEPA | 3 | AGGATTGAGGCCCAAACCGCACCATTT | AAATGGTGCGGTTTGGGCCTCAATCCT |
| C2_ASE1 | *C2* | RABS | 3 | AGGATTGAGGCCCAAAGCGCACCATTT | AAATGGTGCGCTTTGGGCCTCAATCCT |
| C2_ASE2 | *C2* | BEPA | 5 | CATATCTTCTCTGTCTCTGACAACGCC | GGCGTTGTCAGAGACAGAGAAGATATG |
| C2_ASE2 | *C2* | RABS | 5 | CACATCTTCTCTGTCTCTGACAACGCC | GGCGTTGTCAGAGACAGAGAAGATGTG |
| C2_ASE4 | *C2* | BEPA | 13 | TGCTTACCTGCTTGGTCTCAACTCTGC | GCAGAGTTGAGACCAAGCAGGTAAGCA |
| C2_ASE4 | *C2* | RABS | 13 | TGCTTACCTGCTCGGTCTCAACTCTGC | GCAGAGTTGAGACCGAGCAGGTAAGCA |
| C2_ASE6 | *C2* | BEPA | 33 | AAGCAGGTGGAAACAGAGAAGTCTGAG | CTCAGACTTCTCTGTTTCCACCTGCTT |
| C2_ASE6 | *C2* | RABS | 33 | AAGCAAGTGGAAACAGAGAAGTCTGAG | CTCAGACTTCTCTGTTTCCACTTGCTT |
| C3_ASE2 | *C3* | BEPA | 3 | GCCAAAGTCATCAAGAGAGATGGTGGC | GCCACCATCTCTCTTGATGACTTTGGC |

| Set | *MYH3C* target | Allele target | Exon | *k*-mer (forward) | *k*-mer (reverse complement) |
|---|---|---|---|---|---|
| C3_ASE2 | *C3* | RABS | 3 | GCCAAAGTCATCAAGAAAGAGGGTGGC | GCCACCCTCTTTCTTGATGACTTTGGC |
| C3_ASE3 | *C3* | BEPA | 4 | AGGACAGTTAAAGAAGAAGAAATCTTT | AAAGATTTCTTCTTCTTTAACTGTCCT |
| C3_ASE3 | *C3* | RABS | 4 | AGGACAGTTAAAGAAGATGAAATCTTT | AAAGATTTCATCTTCTTTAACTGTCCT |
| C3_ASE4 | *C3* | BEPA | 16 | CCTACCAAGGGCAAGGCTGAGGCCCAC | GTGGGCCTCAGCCTTGCCCTTGGTAGG |
| C3_ASE4 | *C3* | RABS | 16 | CCTACCAAGGGCAAGGCTGAGGGCCAC | GTGGCCCTCAGCCTTGCCCTTGGTAGG |
| C3_ASE5 | *C3* | BEPA | 21 | CACTGGTCACGATGACCCAGGCTTTGT | ACAAAGCCTGGGTCATCGTGACCAGTG |
| C3_ASE5 | *C3* | RABS | 21 | CACTGGTCACGATGACTCAGGCTTTGT | ACAAAGCCTGAGTCATCGTGACCAGTG |
| C3_ASE6 | *C3* | BEPA | 28 | GATGTGCCGTACTCTTGAGGACCAATT | AATTGGTCCTCAAGAGTACGGCACATC |
| C3_ASE6 | *C3* | RABS | 28 | GCTGTGCCGTACTCTTGAGGACCAATT | AATTGGTCCTCAAGAGTACGGCACAGC |
| C3_ASE7 | *C3* | BEPA | 30 | AAACGGTGAGGTGTCTCAGTGGAGATC | GATCTCCACTGAGACACCTCACCGTTT |
| C3_ASE7 | *C3* | RABS | 30 | AAACGGTGAGGTGGCTCAGTGGAGATC | GATCTCCACTGAGCCACCTCACCGTTT |
| C3_ASE10 | *C3* | BEPA | 38 | GAGACTGAGCAGAGACGTGGAGTGGAC | GTCCACTCCACGTCTCTGCTCAGTCTC |
| C3_ASE10 | *C3* | RABS | 38 | GAGGCTGAGCAGAGACGTGGAGTGGAC | GTCCACTCCACGTCTCTGCTCAGCCTC |
| C3_ASE11 | *C3* | BEPA | 38 | GTGGACGCCGTCAAAGGTGTTCGCAAA | TTTGCGAACACCTTTGACGGCGTCCAC |
| C3_ASE11 | *C3* | RABS | 38 | GTGGACGCCGTCAAGGGTGTTCGCAAA | TTTGCGAACACCCTTGACGGCGTCCAC |
| C3_ASE12 | *C3* | BEPA | 40 | ATTGCTGAGTCTCAAGTCAACAAGATG | CATCTTGTTGACTTGAGACTCAGCAAT |
| C3_ASE12 | *C3* | RABS | 40 | ATTGCTGAGTCTCAGGTCAACAAGATG | CATCTTGTTGACCTGAGACTCAGCAAT |

**Table S4. Final K-mers used for RNA-seq analysis, related to Figure 3**

### SUPPLEMENTAL REFERENCES


S1. Roberts Kingman, G.A., Vyas, D.N., Jones, F.C., Brady, S.D., Chen, H.I., Reid, K., Milhaven, M., Bertino, T.S., Aguirre, W.E., Heins, D.C., et al. (2021). Predicting future from past: The genomic basis of recurrent and rapid stickleback evolution. Sci. Adv. *7*, eabg5285. https://doi.org/10.1126/sciadv.abg5285.

S2. Leaver, S.D., and Reimchen, T.E. (2012). Abrupt changes in defence and trophic morphology of the giant threespine stickleback (*Gasterosteus* sp.) following colonization of a vacant habitat. Biol. J. Linn. Soc. *107*, 494–509. https://doi.org/10.1111/j.1095-8312.2012.01969.x.

S3. Jones, F.C., Grabherr, M.G., Chan, Y.F., Russell, P., Mauceli, E., Johnson, J., Swofford, R., Pirun, M., Zody, M.C., White, S., et al. (2012). The genomic basis of adaptive evolution in threespine sticklebacks. Nature *484*, 55–61. https://doi.org/10.1038/nature10944.

S4. Nath, S., Shaw, D.E., and White, M.A. (2021). Improved contiguity of the threespine stickleback genome using long-read sequencing. G3 *11*, jkab007. https://doi.org/10.1093/g3journal/jkab007.

S5. Kingsley, D.M., Zhu, B., Osoegawa, K., De Jong, P.J., Schein, J., Marra, M., Peichel, C., Amemiya, C., Schluter, D., Balabhadra, S., et al. (2004). New genomic tools for molecular studies of evolutionary change in threespine sticklebacks. Behaviour *141*, 1331–1344.

S6. Au, E.H., Weaver, S., Katikaneni, A., Wucherpfennig, J.I., Luo, Y., Mangan, R.J., Wund, M.A., Bell, M.A., and Lowe, C.B. (2025). Genome sequence of a marine threespine stickleback (*Gasterosteus aculeatus*) from Rabbit Slough in the Cook Inlet. G3 *15*, jkaf114. https://doi.org/10.1093/g3journal/jkaf114.

S7. Wucherpfennig, J.I., Howes, T.R., Au, J.N., Au, E.H., Roberts Kingman, G.A., Brady, S.D., Herbert, A.L., Reimchen, T.E., Bell, M.A., Lowe, C.B., et al. (2022). Evolution of stickleback spines through independent *cis*-regulatory changes at *HOXDB*. Nat. Ecol. Evol. *6*, 1537–1552. https://doi.org/10.1038/s41559-022-01855-3.

S8. Peichel, C.L., McCann, S.R., Ross, J.A., Naftaly, A.F.S., Urton, J.R., Cech, J.N., Grimwood, J., Schmutz, J., Myers, R.M., Kingsley, D.M., et al. (2020). Assembly of the threespine stickleback Y chromosome reveals convergent signatures of sex chromosome evolution. Genome Biol. *21*, 177. https://doi.org/10.1186/s13059-020-02097-x.

S9. Ishikawa, A., Kabeya, N., Ikeya, K., Kakioka, R., Cech, J.N., Osada, N., Leal, M.C., Inoue, J., Kume, M., Toyoda, A., et al. (2019). A key metabolic gene for recurrent freshwater colonization and radiation in fishes. Science *364*, 886–889. https://doi.org/10.1126/science.aau5656.

S10. Thorburn, D.-M.J., Sagonas, K., Binzer-Panchal, M., Chain, F.J.J., Feulner, P.G.D., Bornberg-Bauer, E., Reusch, T.B.H., Samonte-Padilla, I.E., Milinski, M., Lenz, T.L., et al. (2023). Origin matters: Using a local reference genome improves measures in population genomics. Mol. Ecol. Resour. *23*, 1706–1723. https://doi.org/10.1111/1755-0998.13838.

S11. Adhikari, D., Karlsen, B.O., Jørgensen, T.E., Johansen, S.D., Nordeide, J.T., and Moum, T.B. (2025). The genomics of postglacial vicariance and freshwater adaptations in European subarctic threespine sticklebacks. Front. Ecol. Evol. *13*, 1546874. https://doi.org/10.3389/fevo.2025.1546874.
